# Changes in brain connectivity and neurovascular dynamics during dexmedetomidine-induced loss of consciousness

**DOI:** 10.1101/2024.10.04.616650

**Authors:** Panagiotis Fotiadis, Andrew R. McKinstry-Wu, Sarah M. Weinstein, Philip A. Cook, Mark Elliott, Matthew Cieslak, Jeffrey T. Duda, Theodore D. Satterthwaite, Russell T. Shinohara, Alexander Proekt, Max B. Kelz, John A. Detre, Dani S. Bassett

## Abstract

Understanding the neurophysiological changes that occur during loss and recovery of consciousness is a fundamental aim in neuroscience and has marked clinical relevance. Here, we utilize multimodal magnetic resonance neuroimaging to investigate changes in regional network connectivity and neurovascular dynamics as the brain transitions from wakefulness to dexmedetomidine-induced unconsciousness, and finally into early-stage recovery of consciousness. We observed widespread decreases in functional connectivity strength across the whole brain, and targeted increases in structure-function coupling (SFC) across select networks— especially the cerebellum—as individuals transitioned from wakefulness to hypnosis. We also observed robust decreases in cerebral blood flow (CBF) across the whole brain—especially within the brainstem, thalamus, and cerebellum. Moreover, hypnosis was characterized by significant increases in the amplitude of low-frequency fluctuations (ALFF) of the resting-state blood oxygen level-dependent signal, localized within visual and somatomotor regions. Critically, when transitioning from hypnosis to the early stages of recovery, functional connectivity strength and SFC—but not CBF—started reverting towards their awake levels, even before behavioral arousal. By further testing for a relationship between connectivity and neurovascular alterations, we observed that during wakefulness, brain regions with higher ALFF displayed lower functional connectivity with the rest of the brain. During hypnosis, brain regions with higher ALFF displayed weaker coupling between structural and functional connectivity. Correspondingly, brain regions with stronger functional connectivity strength during wakefulness showed greater reductions in CBF with the onset of hypnosis. Earlier recovery of consciousness was associated with higher baseline (awake) levels of functional connectivity strength, CBF, and ALFF, as well as female sex. Across our findings, we also highlight the role of the cerebellum as a recurrent marker of connectivity and neurovascular changes between states of consciousness. Collectively, these results demonstrate that induction of, and emergence from dexmedetomidine-induced unconsciousness are characterized by widespread changes in connectivity and neurovascular dynamics.

## INTRODUCTION

Unconsciousness is a multifaceted state; it can occur in a naturalistic setting such as during sleep, in response to sedative/hypnotic drugs, or after a neurological event such as stroke, traumatic brain injury, or epileptic seizure. In addition to its clinical implications, understanding the neural, hemodynamic, and metabolic events taking place during loss of consciousness may provide further insights into the neural basis of consciousness. Anesthesia has emerged as an effective tool in this endeavor as it allows for the reversible manipulation of wakefulness in a highly controlled setting, under a variety of pharmacological agents.

One such agent, dexmedetomidine, has been particularly versatile in the clinical and research setting, due to its ability to induce hypnosis while still allowing for the easy awakening of the participant using gentle tactile or verbal stimulation.^1,2^ It is typically used as a sedative/hypnotic and analgesic agent in the intensive care unit as well as during various medical and surgical procedures.^3,4^ In the research setting, it has been used to pharmacologically induce a state that neurophysiologically resembles non-rapid eye movement (NREM) sleep.^5–8^ An α_2_-adrenergic receptor agonist, dexmedetomidine selectively binds to pre-synaptic a_2_-adrenergic receptors on neurons that project to major arousal centers across the brain, leading to a decrease in their firing rate and hence norepinephrine release.^6,9–11^ Dexmedetomidine is also thought to induce hypnotic effects by engaging endogenous NREM sleep-promoting neural networks.^7,10,12,13^

Dexmedetomidine-induced loss of consciousness is accompanied by a cascade of neurophysiological changes spanning cortical and subcortical brain regions. Using tools from graph theory, prior work has shown that the dexmedetomidine-induced unconscious state is characterized by a decreased capacity for efficient information transmission, both locally and globally.^14^ These changes result from brain regions reducing their overall synchronization strength with other regions, a feature that is particularly evident within strongly connected areas.^14^ Interestingly, cortico-cortical functional connectivity within two higher-order association networks—the default mode network and the fronto-parietal network—defined using resting-state functional magnetic resonance imaging (fMRI) was found to be preserved during the unconscious state; functional connectivity between the thalamus and higher-order association networks such as the default mode, executive control, and salience networks, however, was significantly disrupted.^12,15^

Previous work has also shown that dexmedetomidine affects regional cerebral blood flow (CBF) and metabolism, preferentially decreasing CBF and glucose metabolism in the thalamus, default mode, and bilateral fronto-parietal networks.^12^ Recovery from dexmedetomidine-induced unconsciousness has been associated with (i) a sustained reduction in CBF,^12^ (ii) restored thalamic functional connectivity to the default mode network,^12^ and (iii) increased functional activation of the hypothalamus, thalamus, brainstem, cerebellum, anterior cingulate cortex, and portions of the lateral orbital frontal and parietal lobes.^2^ These findings underscore how interactions between posterior circulation regions and anterior regions give rise to volitional consciousness.^1,2^

Changes in blood flow have been known to accompany changes in neural activity—a process referred to as neurovascular coupling. By quantifying this relationship, recent work suggests a dynamical coupling between CBF and the amplitude of low frequency (0.01–0.08 Hz) fluctuations (ALFF) in the blood oxygen level-dependent (BOLD) signal.^16–19^ The power of low-frequency fluctuations has also been reported to change during anesthetic induction. Indeed, anesthetic agents including dexmedetomidine,^20,21^ propofol,^21–24^ sevoflurane,^22,25^ and high-dose ketamine,^22,26,27^ among others,^25,28,29^ have all been reported to induce an increase in the power of slow/delta frequencies (0.1–4 Hz) in humans, based on electroencephalogram (EEG) recordings. Increases in slow wave activity (< 1 Hz) are also evident during NREM sleep^24,30^ and coma (vegetative state).^31,32^ These findings collectively highlight the role of low frequency oscillations in electrophysiologically characterizing unconsciousness (although see Refs.^33–35^ for interesting case-studies wherein behavioral levels of consciousness were dissociated from EEG features such as slow wave oscillations).

Building upon this work, we utilized multimodal non-invasive magnetic resonance neuroimaging to further characterize the regional connectivity and neurovascular alterations that occur in the human brain as it transitions from the awake state to a state of dexmedetomidine-induced unconsciousness and finally to the early-stage recovery of consciousness. To that end, we assessed two connectivity markers: functional connectivity strength and structure-function coupling (SFC)—a metric^36^ that quantifies the correlation between a region’s patterns of functional and structural connectivity to the rest of the brain—and two neurovascular markers: CBF and ALFF. We examined functional connectivity strength, SFC, and CBF using arterial spin labeling (ASL) imaging acquired across three awake time-points, two hypnotic time-points, and one recovery time-point, and ALFF using resting-state BOLD fMRI acquired across one awake and one hypnotic time-point (**Figure 1**).

**Figure 1.**
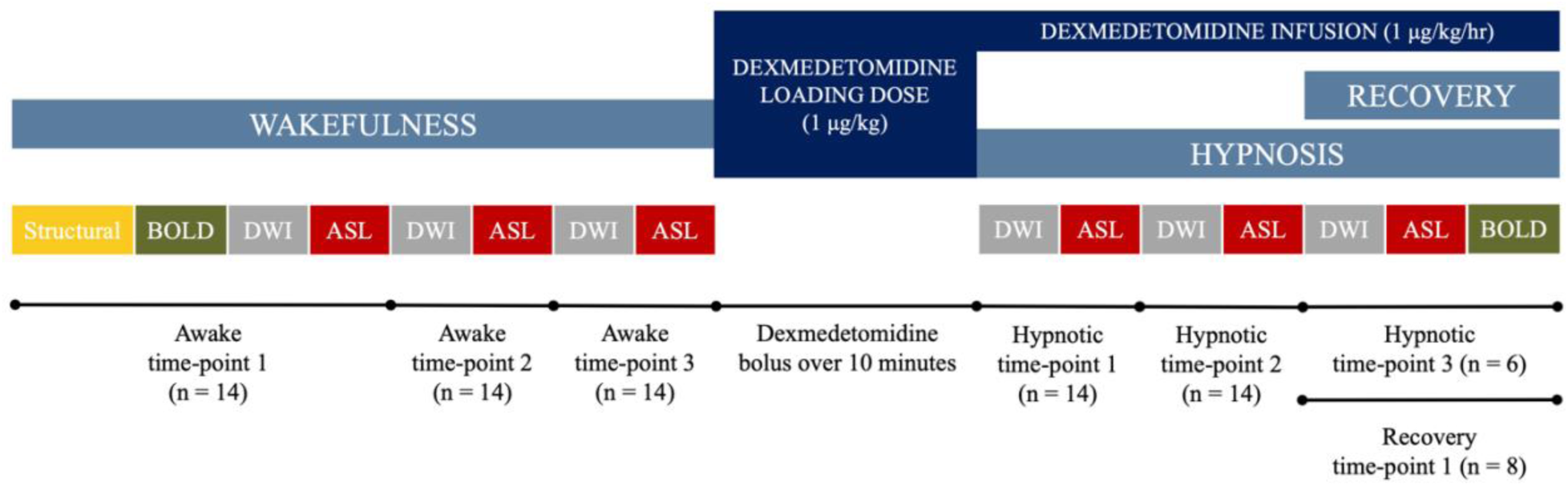
Experimental design. Schematic illustration of the experimental setup used in this study. Healthy individuals (n = 14) underwent a magnetic resonance imaging (MRI) battery consisting of structural (T1-weighted), resting-state functional MR, diffusion-weighted, and arterial spin labeling imaging, while they were awake (awake state). Each awake time-point is denoted, followed by the total number of individuals who completed it. Dexmedetomidine was loaded over 10 minutes and then administered as a continuous infusion for the duration of the “hypnosis” scans. Once individuals were behaviorally unconscious, the same MRI battery was repeated during hypnosis (hypnotic state). For hypnotic time-points 1 and 2, data from all individuals (n = 14) were used; for hypnotic time-point 3, only data from a subset of individuals who remained unconscious were used (n = 6), as the rest of the sample had already started recovering from the hypnosis. This latter subset (n = 8) was labeled as the “recovery sample;” data from these individuals were used to define recovery time-point 1. BOLD: blood oxygen level-dependent signal; DWI: diffusion-weighted imaging; ASL: arterial spin labeling.

Given the aforementioned decreased capacity for efficient information transmission during dexmedetomidine-induced loss of consciousness, we first asked whether functional connectivity strength and SFC respectively decrease and increase during the transition from wakefulness to hypnosis, and reverse towards recovery. Moreover, given the reported decreases in CBF and glucose metabolism across select brain regions during loss of consciousness, we next asked whether CBF decreases across the brain during pharmacological hypnosis. Given the dynamic relationship between blood flow and low frequency fluctuations, as well as the previously reported increases in the power of low frequency fluctuations in EEG during various states of unconsciousness, we then asked whether ALFF also increases during dexmedetomidine-induced hypnosis. To assess whether dexmedetomidine-induced changes in connectivity and neurovascular dynamics are site-specific, we tested these hypotheses across cerebellar, subcortical, and cortical networks and regions spanning the sensory-association axis of cortical organization, ranging from primary sensory and motor cortices to higher-order association cortices such as the fronto-parietal and default mode networks.^37–39^ Lastly, we investigated whether there is a relationship between changes in connectivity and neurovascular dynamics as the brain loses and recovers consciousness.

## RESULTS

### Connectivity changes from wakefulness to hypnosis

We first examined how functional connectivity strength—estimated via ASL imaging (see **Methods**)—and structure-function coupling (SFC) change from wakefulness to dexmedetomidine-induced unconsciousness, at three levels of granularity: the whole-brain level, the network level, and the regional level. Functional connectivity strength and SFC were defined using ASL imaging,^40,41^ as it had been acquired across 3 awake time-points, 2 hypnotic time-points, and one recovery time-point (**Figure 1**).

Functional connectivity strength remained similar across the 3 awake time-points, and separately across the 2 hypnotic time-points. During the transition from the 3^rd^ awake time-point to the 1^st^ hypnotic time-point, we observed a significant decrease in overall functional connectivity strength (Repeated measures [rm]ANOVA; Tukey’s post-hoc pairwise comparisons: 6% decrease; adjusted *p_HSD_* = 0; **Figure 2A**). An overall decrease in functional connectivity strength from the average awake to the average hypnotic state was also observed across brain regions (5.1% decrease; paired samples Wilcoxon test: *p* = 2.1 x 10^-23^; **Figure 2B**) and across individuals (5% decrease; two-sided paired *t*-test; *p* = 0.008; **Figure 3A**).

**Figure 2.**
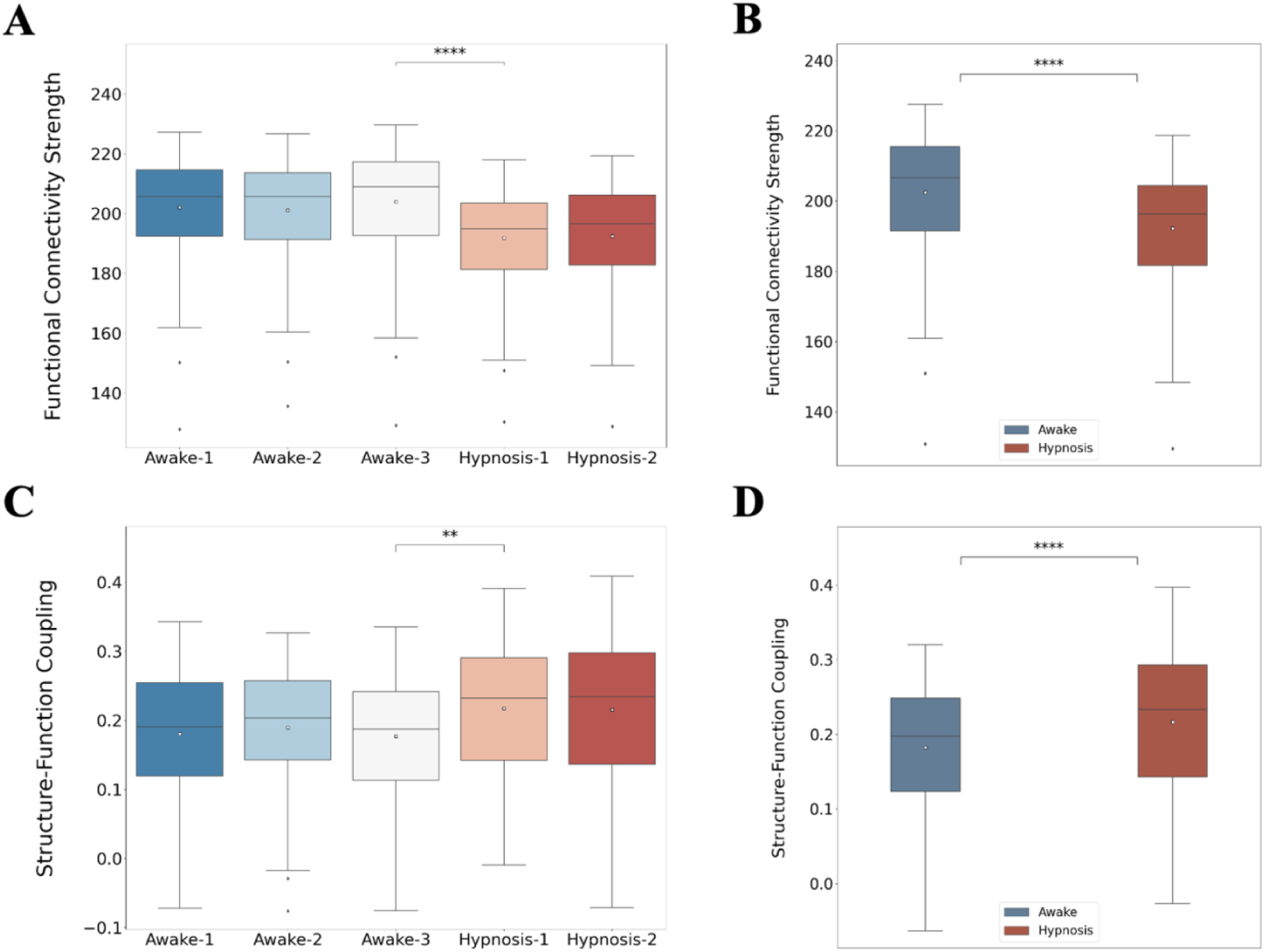
Whole-brain changes in functional connectivity strength and structure-function coupling from wakefulness to dexmedetomidine-induced unconsciousness. **A**: Changes in whole-brain functional connectivity strength across the 3 awake and 2 hypnotic time-points (n = 132 brain regions/datapoints for each boxplot, averaged across all individuals). **B**: Changes in whole-brain functional connectivity strength across the average awake state and the average hypnotic state (n = 132 brain regions/datapoints for each boxplot, averaged across all individuals). **C**: Changes in whole-brain structure-function coupling across the 3 awake and 2 hypnotic time-points (n = 132 brain regions/datapoints for each boxplot, averaged across all individuals). **D**: Changes in whole-brain structure-function coupling across the average awake state and the average hypnotic state (n = 132 brain regions/datapoints for each boxplot, averaged across all individuals). *p*-value annotation legend: non-significant (ns): 0.05 < *p* ≤ 1, *: 0.01 < *p* ≤ 0.05, **: 0.001< *p* ≤ 0.01, ***: 10^-4^ < *p* ≤ 10^-3^, ****: *p* ≤ 10^-4^; *p*-values in Figures 2A and **2C** correspond to repeated measures ANOVA tests followed by *post-hoc* correction for multiple comparisons (Tukey’s honestly significant difference [HSD] test), while *p*-values in Figures 2B and **2D** correspond to paired samples Wilcoxon tests.

**Figure 3.**
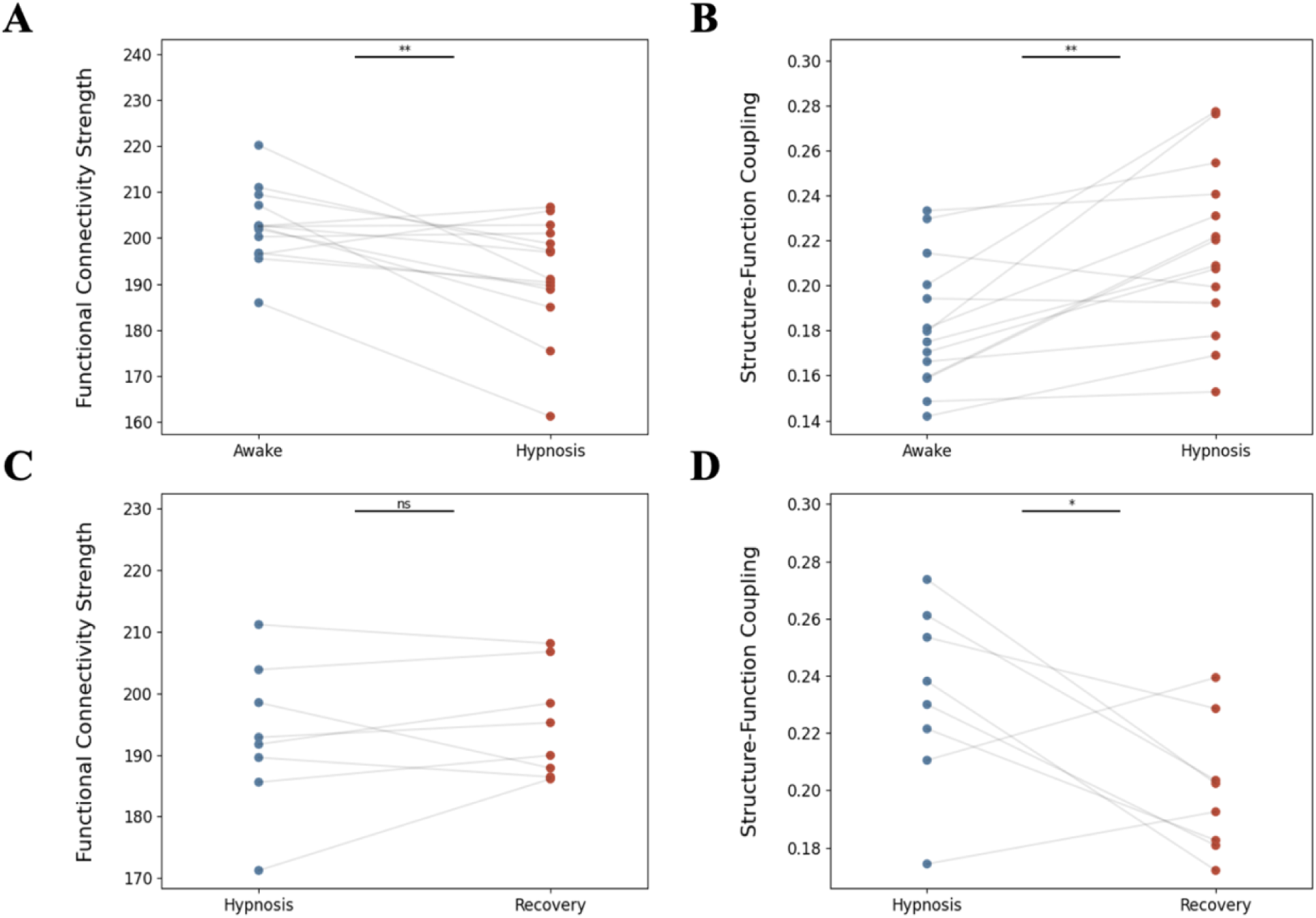
Whole-brain changes in connectivity variables from dexmedetomidine-induced unconsciousness to early-stage recovery. **A**: Changes in whole-brain functional connectivity strength for each individual across the average awake state and the average hypnotic state (n = 14 datapoints for each state, denoting each individual). **B**: Changes in whole-brain structure-function coupling for each individual across the average awake state and the average hypnotic state (n = 14 datapoints for each state, denoting each individual). **C**: Changes in whole-brain functional connectivity strength for each individual across the last hypnotic time-point and the recovery time-point (n = 8 datapoints for each state, denoting each individual that recovered consciousness before the end of the experiment). **D**: Changes in whole-brain structure-function coupling for each individual across the last hypnotic time-point and the recovery time-point (n = 8 datapoints for each state, denoting each individual that recovered consciousness before the end of the experiment). *p*-value annotation legend: non-significant (ns): 0.05 < *p* ≤ 1, *: 0.01 < *p* ≤ 0.05, **: 0.001< *p* ≤ 0.01, ***: 10^-4^ < *p* ≤ 10^-3^, ****: *p* ≤ 10^-4^; *p*-values here correspond to paired *t*-tests.

Narrowing our focus from the whole-brain level to the network level, we focused on previously established^37^ resting-state functional networks spanning the sensory-association axis of the human brain: the visual, somatomotor, dorsal attention, ventral attention, limbic, fronto-parietal, default mode, subcortical, and cerebellar networks. Changes in functional connectivity strength were starkly evident across all networks, with each network decreasing its strength as it entered hypnosis (**Supplementary Table 1**; **Supplementary Figure 1**). Out of all networks considered, the visual and somatomotor networks displayed the largest percent decreases in magnitude (visual: 9.1% decrease; paired samples Wilcoxon test with Benjamini-Hochberg correction for multiple comparisons: *p_BH_* = 3.4 x 10^-5^, somatomotor: 6% decrease; *p_BH_* = 2.3 x 10^-5^).

Concentrating our focus even further, we next examined how individual brain regions altered their functional connectivity strength during the shift from wakefulness to hypnosis. Multiple regions significantly decreased their functional connectivity strength, including 3 cerebellar sub-regions, 2 subcortical grey nuclei (left putamen and left hippocampus), and cortical regions spanning across the visual (16 regions), somatomotor (13 regions), dorsal attention (5 regions), ventral attention (3 regions), fronto-parietal (3 regions), and default mode (9 regions) networks (**Supplementary Table 4**). Among these regions, functional connectivity strength decreases were most prominent within occipital and temporal regions involved in visual and auditory processing, including the cuneal cortex (an average 12.5% decrease across individuals; two-sided paired *t*-test followed by Benjamini-Hochberg correction for multiple comparisons: *p_BH_* = 2.7 x 10^-4^), intracalcarine cortex (an average 11.1% decrease across individuals; *p_BH_* = 7.3 x 10^-4^), supracalcarine cortex (an average 11.5% decrease across individuals; *p_BH_* = 4 x 10^-4^), and lingual gyrus (an average 10.3% decrease across individuals; *p_BH_* = 3 x 10^-4^).

We then assessed how the relationship between a region’s functional connectivity and underlying structural connectivity changes as the human brain transitions from wakefulness to dexmedetomidine-induced unconsciousness. For that purpose, we turned to SFC. Although similar in magnitude across the 3 awake time-points and across the 2 hypnotic time-points, SFC significantly increased during the transition from the last awake time-point to the first hypnotic time-point (rmANOVA; Tukey’s post-hoc pairwise comparisons: 22.8% increase; adjusted *p_HSD_* = 0.003; **Figure 2C**). Furthermore, the average hypnotic state was characterized by higher SFC than the average awake state across brain regions (18.7% increase; paired samples Wilcoxon test: *p* = 1.5 x 10^-13^; **Figure 2D**) and across individuals (19.5% increase; two-sided paired *t*-test; *p* = 0.002; **Figure 3B**).

On the network level, SFC increased in magnitude in unimodal visual and motor cortices (visual: 18% increase; paired samples Wilcoxon test with Benjamini-Hochberg correction for multiple comparisons: *p_BH_* = 2.1 x 10^-4^, somatomotor: 10.1% increase; *p_BH_* = 0.013), higher-order transmodal association cortices (fronto-parietal: 15.1% increase; *p_BH_* = 0.028, default mode: 13.7% increase; *p_BH_* = 0.002), and cerebellar networks (51.6% increase; *p_BH_* = 2.7 x 10^-7^) (**Supplementary Figure 2**).

On the regional level, SFC increased in one subcortical structure (right pallidum) and 3 cortical regions located in the visual (left lingual), fronto-parietal (right frontal pole), and default mode (right paracingulate gyrus) networks, and decreased in one cortical region located in the limbic network (right posterior inferior temporal gyrus) (**Supplementary Table 5**). The most pronounced increases in SFC, however, were localized within the cerebellum, and specifically across 7 cerebellar sub-regions (average 134.1% increase in SFC in these areas, across individuals; *p_BH_* = 0.023), with certain sub-regions increasing their SFC more than three-fold (**Supplementary Table 5**).

### Connectivity changes from hypnosis to early-stage recovery

Next, we examined how functional connectivity strength and SFC changed during the transition from hypnosis to early-stage recovery (i.e., between the 2^nd^ hypnotic and 1^st^ recovery time-points—see **Figure 1**) on a subset of the population that started regaining consciousness towards the end of the hypnosis protocol, while still receiving a steady infusion of dexmedetomidine (**Figure 1**).

Although there was no change in overall functional connectivity strength across individuals (1.1% increase; two-sided paired *t*-test; *p* = 0.527; **Figure 3C**), there was a subtle yet significant increase in functional connectivity strength across brain regions, after averaging across individuals, (0.9% increase; paired samples Wilcoxon test: *p* = 2.2 x 10^-5^) towards awake levels. Increases in functional connectivity strength were localized within the visual (3.9% increase; paired samples Wilcoxon test with Benjamini-Hochberg correction for multiple comparisons: *p_BH_* = 6.9 x 10^-4^) and default mode (1.9% increase; *p_BH_* = 5.7 x 10^-4^) networks, among a few other networks (**Supplementary Table 2**; **Supplementary Figure 3**).

Moreover, decreases in whole-brain SFC towards the early-stage recovery time-point were pronounced across brain regions (14% decrease; paired samples Wilcoxon test: *p* = 5.6 x 10^-13^) and across individuals (12.6% decrease; two-sided paired *t*-test; *p* = 0.045; **Figure 3D**), with SFC magnitude reverting back towards awake levels. These decreases were most prominently expressed within the cerebellar network (23.4% decrease; paired samples Wilcoxon test with Benjamini-Hochberg correction for multiple comparisons: *p_BH_* = 1.9 x 10^-5^; **Supplementary Figure 4**); other networks that significantly increased their SFC included the somatomotor, ventral attention, fronto-parietal, default mode, and subcortical networks (**Supplementary Table 3**).

Narrowing our focus once again from the network level to the individual brain region level, there were no specific brain regions whose functional connectivity strength or SFC changed significantly during the transition from hypnosis to early-stage recovery, after correction for multiple comparisons.

### Neurovascular changes from wakefulness to hypnosis

To investigate the neurovascular changes that take place as the brain transitions from wakefulness to dexmedetomidine-induced unconsciousness, we next focus on two markers: the amplitude of low-frequency fluctuations (ALFF; 0.01–0.08 Hz) within the resting-state BOLD signal, and the CBF obtained from ASL imaging. As with the connectivity analyses, we address this question at the whole-brain, network, and regional levels.

On average, brain regions’ ALFF increased during hypnosis (25.8% increase; paired samples Wilcoxon test: *p* = 2.2 x 10^-16^; **Figure 4A**). This result was recapitulated across individuals (27% increase; two-sided paired *t*-test: *p* = 0.002; **Figure 4C**). Increases in ALFF were localized across multiple networks (**Supplementary Figure 5**), including the unimodal primary visual and motor cortices (visual: 50.5% increase; paired samples Wilcoxon test with Benjamini-Hochberg correction for multiple comparisons: *p_BH_* = 4.6 x 10^-5^, somatomotor: 54.6% increase; *p_BH_* = 3.4 x 10^-5^), the dorsal attention network (29.6% increase; *p_BH_* = 0.028), as well as the higher-order default mode (31.7% increase; *p_BH_* = 1.7 x 10^-5^) and limbic (19.4% increase; *p_BH_* = 0.001) networks. After dividing each brain region’s ALFF signal by its overall BOLD signal power—a metric commonly referred to as fractional ALFF^42^—the directionality of results inverted, with fractional ALFF decreasing from wakefulness to hypnosis across brain regions (10.7% decrease; paired samples Wilcoxon test: *p* = 1.3 x 10^-13^) and individuals (10.4% decrease; two-sided paired *t*-test: *p* = 0.033; **Supplementary Figure 6**). As with ALFF, these decreases were especially localized within the unimodal primary visual and motor cortices (visual: 26.3% decrease; paired samples Wilcoxon test with Benjamini-Hochberg correction for multiple comparisons: *p_BH_* = 1.4 x 10^-4^, somatomotor: 16.1% decrease; *p_BH_* = 6.9 x 10^-5^; **Supplementary Figure 7**).

**Figure 4.**
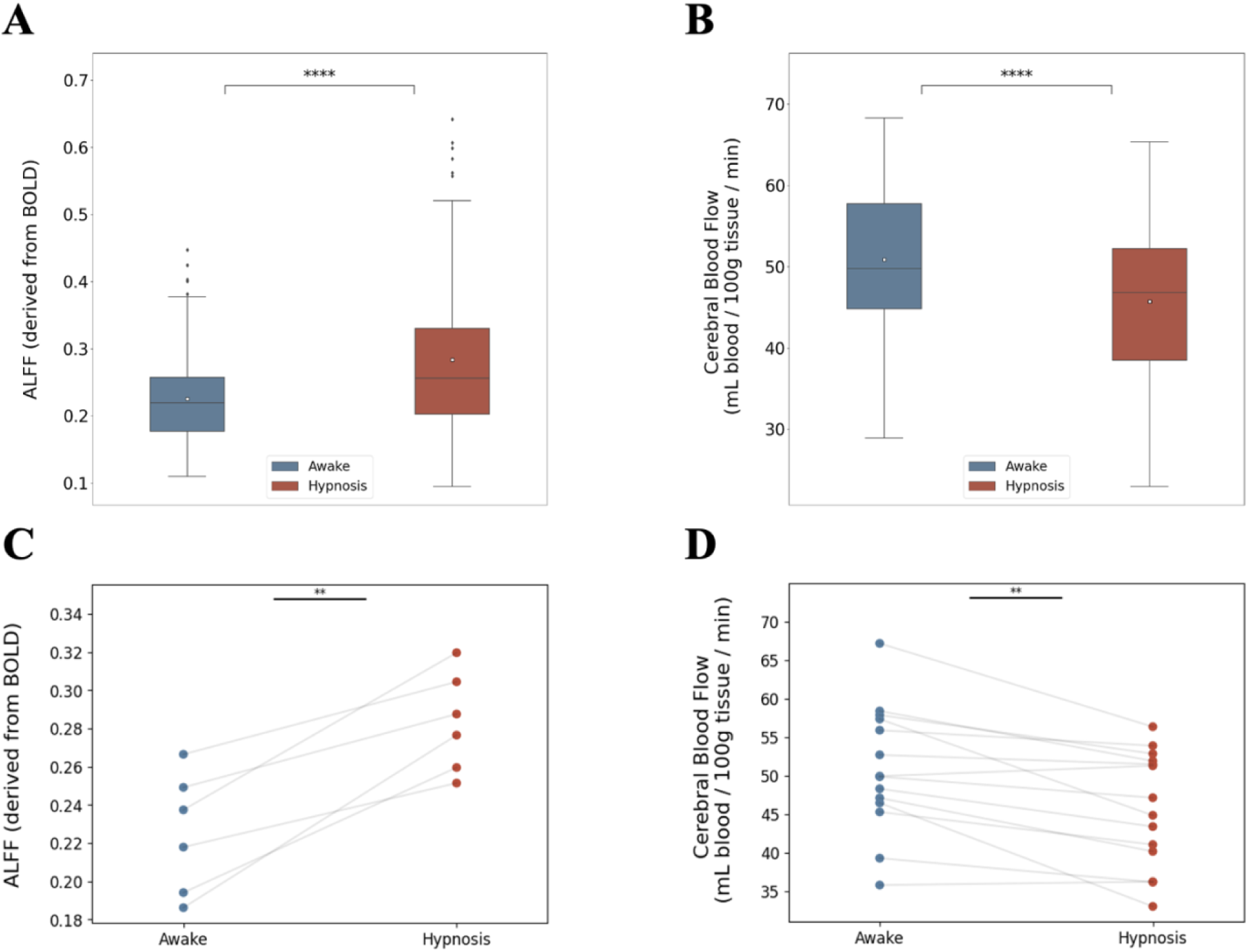
Whole-brain changes in amplitude of low-frequency fluctuations and cerebral blood flow from wakefulness to dexmedetomidine-induced unconsciousness. **A**: Changes in whole-brain amplitude of low-frequency fluctuations (ALFF) across the average awake state and the average hypnotic state (n = 132 brain regions/datapoints for each boxplot, averaged across all individuals). **B**: Changes in whole-brain cerebral blood flow (CBF) across the average awake state and the average hypnotic state (n = 132 brain regions/datapoints for each boxplot, averaged across all individuals). **C**: Changes in whole-brain ALFF for each individual across the average awake state and the average hypnotic state (n = 6 datapoints for each state, denoting each individual that remained unconscious until the end of the experiment). **D**: Changes in whole-brain CBF for each individual across the average awake state and the average hypnotic state (n = 14 datapoints for each state, denoting each individual). *p*-value annotation legend: non-significant (ns): 0.05 < *p* ≤ 1, *: 0.01 < *p* ≤ 0.05, **: 0.001< *p* ≤ 0.01, ***: 10^-4^ < *p* ≤ 10^-3^, ****: *p* ≤ 10^-4^; *p*-values in Figures 4A and **4B** correspond to paired samples Wilcoxon tests, while *p*-values in Figures 4C and **4D** correspond to paired *t*-tests.

On the regional level, ALFF significantly increased across brain regions within the visual (9 regions), somatomotor (4 regions), dorsal attention (1 region), and limbic (1 region) networks (**Supplementary Table 7**). Among these regions, ALFF increases were particularly robust within occipital and temporal regions involved in visual, somatomotor, and auditory processing, including the temporal occipital fusiform cortex (an average 78.1% increase across individuals; two-sided paired *t*-test followed by Benjamini-Hochberg correction for multiple comparisons: *p_BH_* = 0.031), left planum temporale (an average 121% increase across individuals; *p_BH_* = 0.031), and left Heschl’s gyrus (an average 100.7% increase across individuals; *p_BH_* = 0.031). Similarly, fractional ALFF decreased across these regions during hypnosis (**Supplementary Table 8**).

Following the opposite directionality than ALFF, CBF decreased, overall, during the transition from wakefulness to hypnosis across brain regions (10.1% decrease; paired samples Wilcoxon test: *p* = 2.1 x 10^-23^; **Figure 4B**) and across individuals (9.7% decrease; two-sided paired *t*-test; *p* = 0.001; **Figure 4D**). When examined across the different awake and hypnotic time-points, CBF remained similar across the 3 awake time-points, and separately across the 2 hypnotic time-points. During the transition from the 3^rd^ awake time-point to the 1^st^ hypnotic time-point, however, there was a significant drop in CBF across brain regions (rmANOVA; Tukey’s post-hoc pairwise comparisons: 7.8% decrease; adjusted *p_HSD_* = 0.002; **Figure 5A**).

**Figure 5.**
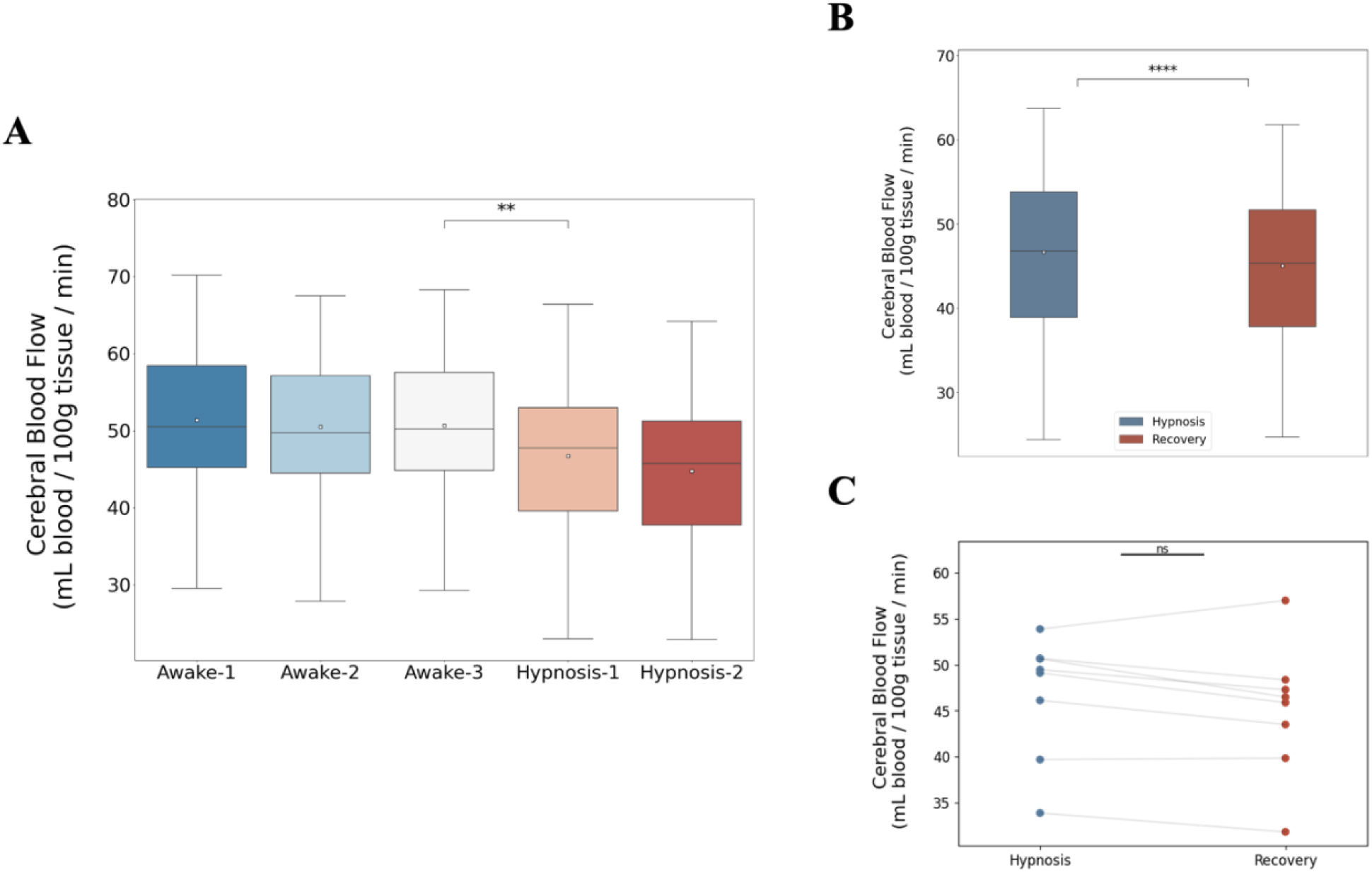
Whole-brain changes in cerebral blood flow from wakefulness to dexmedetomidine-induced unconsciousness. **A**: Changes in whole-brain cerebral blood flow across the 3 awake and 2 hypnotic time-points (n = 132 brain regions/datapoints for each boxplot, averaged across all individuals). **B**: Changes in whole-brain cerebral blood flow across the last hypnotic time-point and the recovery time-point (n = 132 brain regions/datapoints for each boxplot, averaged across all individuals). **C**: Changes in whole-brain CBF for each individual across the last hypnotic time-point and the recovery time-point (n = 8 datapoints for each state, denoting each individual that recovered consciousness before the end of the experiment). *p*-value annotation legend: non-significant (ns): 0.05 < *p* ≤ 1, *: 0.01 < *p* ≤ 0.05, **: 0.001< *p* ≤ 0.01, ***: 10^-4^ < *p* ≤ 10^-3^, ****: *p* ≤ 10^-4^; the *p*-value in Figure 5A corresponds to a repeated measures ANOVA test followed by *post-hoc* correction for multiple comparisons (Tukey’s honestly significant difference [HSD] test), the *p*-value in Figure 5B corresponds to a paired samples Wilcoxon test, and the *p*-value in Figure 5C corresponds to a paired *t*-test.

Changes in CBF were ubiquitous across the brain, with all networks diminishing their CBF magnitude during the transition from wakefulness to dexmedetomidine-induced unconsciousness (**Supplementary Table 6**; **Supplementary Figure 8**). Networks that displayed particularly robust drops in CBF included the visual (11.1% decrease; paired samples Wilcoxon test with Benjamini-Hochberg correction for multiple comparisons: *p_BH_* = 3.4 x 10^-5^), subcortical (11.4% decrease; *p_BH_* = 9.2 x 10^-5^), and cerebellar (17.9% decrease; *p_BH_* = 2.7 x 10^-7^) networks.

On the regional level, the majority of regions considered (80 out of the total of 132 regions; **Supplementary Table 9**) showed significant decreases in their CBF during the transition to hypnosis. Among these regions, the thalamus (an average 18% decrease across individuals; *p_BH_* = 6.7 x 10^-4^), brainstem (an average 19.5% decrease across individuals; *p_BH_* = 5.8 x 10^-4^), and cerebellum (an average 17.4% decrease across individuals; *p_BH_* = 0.016) displayed the largest decreases in magnitude (**Figure 6**; **Supplementary Table 9**).

**Figure 6.**
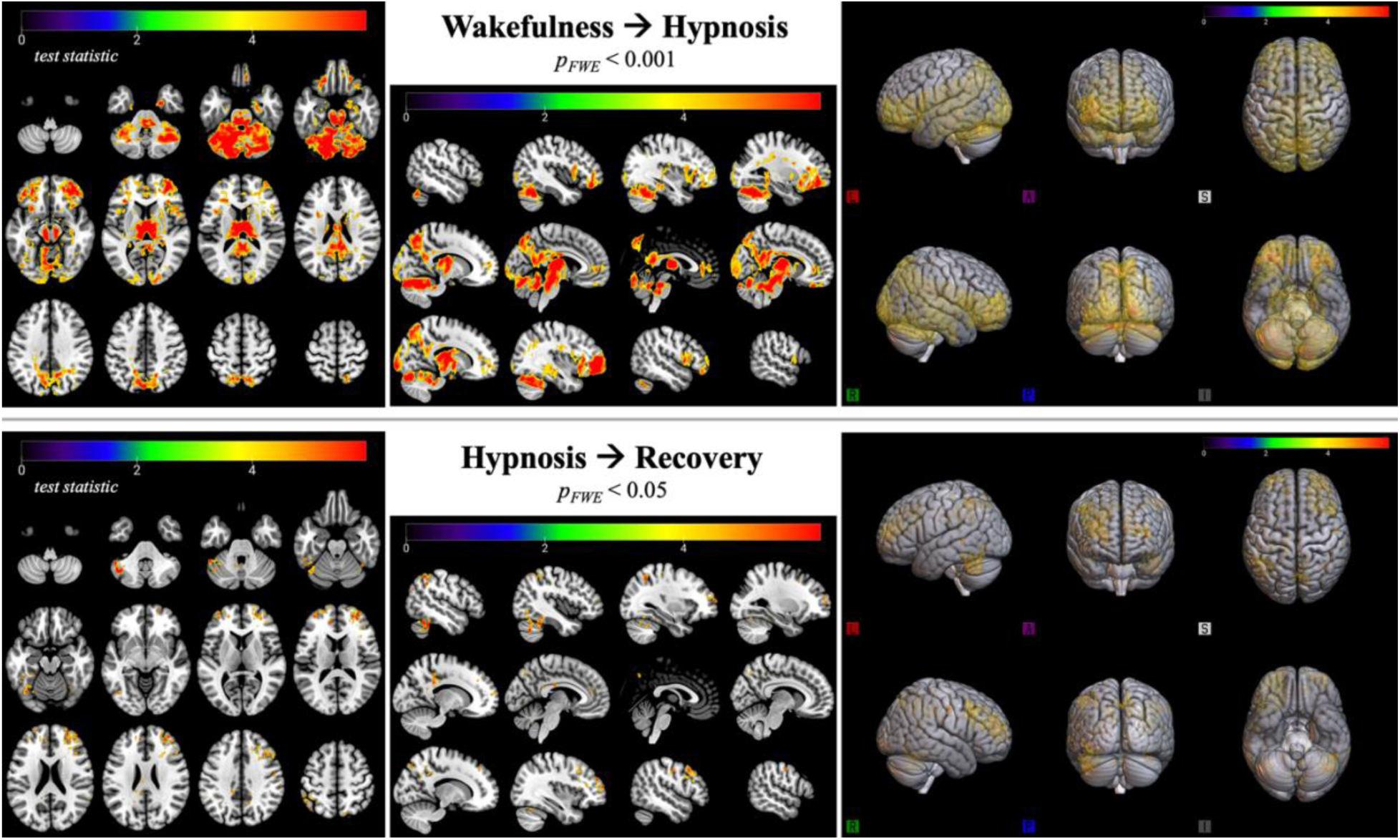
Regional changes in cerebral blood flow during the transition from wakefulness to dexmedetomidine-induced unconsciousness, and finally to early-stage recovery. Top row: Cerebral blood flow (CBF) changes during the transition from wakefulness to hypnosis along axial brain slices (left column), sagittal slices (middle column), and on a 3D rendered standardized (MNI152) brain (right column), across all individuals (n = 14). Warm colors indicate brain regions where CBF is higher during wakefulness, compared to hypnosis. **Bottom row:** CBF changes during the transition from hypnosis to early-stage recovery of consciousness along axial brain slices (left column), sagittal slices (middle column), and on a 3D rendered standardized (MNI152) brain (right column), across individuals within the recovery sample (n = 8). Warm colors indicate brain regions where CBF is higher during hypnosis, compared to recovery. All spatial maps were generated using FSL’s *randomise* tool, a non-parametric permutation inference tool (number of permutations for each analysis = 5,000). *test statistic*: threshold-free cluster enhancement-based statistic^110^ derived from FSL’s *randomise* tool; *p_FWE_*: p-value corrected for family-wise error.

### Neurovascular changes from hypnosis to early-stage recovery

Interestingly, instead of reverting back to awake levels, whole-brain CBF continued decreasing during the transition from hypnosis to early-stage recovery, across brain regions (3.6% decrease; paired samples Wilcoxon test: *p* = 1.9 x 10^-19^; **Figure 5B**). This overall sustained decrease in CBF persisted across all networks examined (**Supplementary Figure 9**). BOLD signal-derived ALFF could not be compared between the states of hypnosis and early-stage recovery across the same individuals, as it was only acquired at the beginning and end of the experiment (**Figure 1**; **Supplemental Material: Methodological Limitations**).

In the sub-sample of individuals who started recovering consciousness before the end of the hypnosis protocol, there were no brain regions that showed significant changes in their CBF levels during the transition from hypnosis to early-stage recovery.

### Individuals who remained unconscious versus individuals who regained consciousness

We next examined whether the aforementioned connectivity and neurovascular markers could distinguish between individuals in our sample who unexpectedly regained consciousness before the end of the experimental session (recovery sample: n = 8) and those who remained unconscious (unconscious sample: n = 6). To that end, we compared the two samples’ connectivity (functional connectivity strength and SFC) and neurovascular metrics (ALFF and CBF) during wakefulness, before the onset of hypnosis. It is of note that all individuals within the recovery sample were female (see **Methods**).

In network connectivity, individuals within the recovery sample had significantly higher overall functional connectivity strength (percentage difference: 4.1%; two-sided Mann-Whitney U test: *p* = 3 x 10^-4^; **Figure 7A**) while awake, compared to the individuals who remained unconscious. These increases were evident across the visual (percentage difference: 4.3%; uncorrected *p* = 0.016 [*p*_BH_ = 0.072, after correction for multiple comparisons across the 9 networks examined]), default mode (percentage difference: 2.1%; uncorrected *p* = 0.012 [*p_BH_* = 0.072, after correction for multiple comparisons]), and cerebellar (percentage difference: 7.8%; uncorrected *p* = 0.032 [*p_BH_* = 0.095, after correction for multiple comparisons]) networks (**Supplementary Figure 10**).

**Figure 7.**
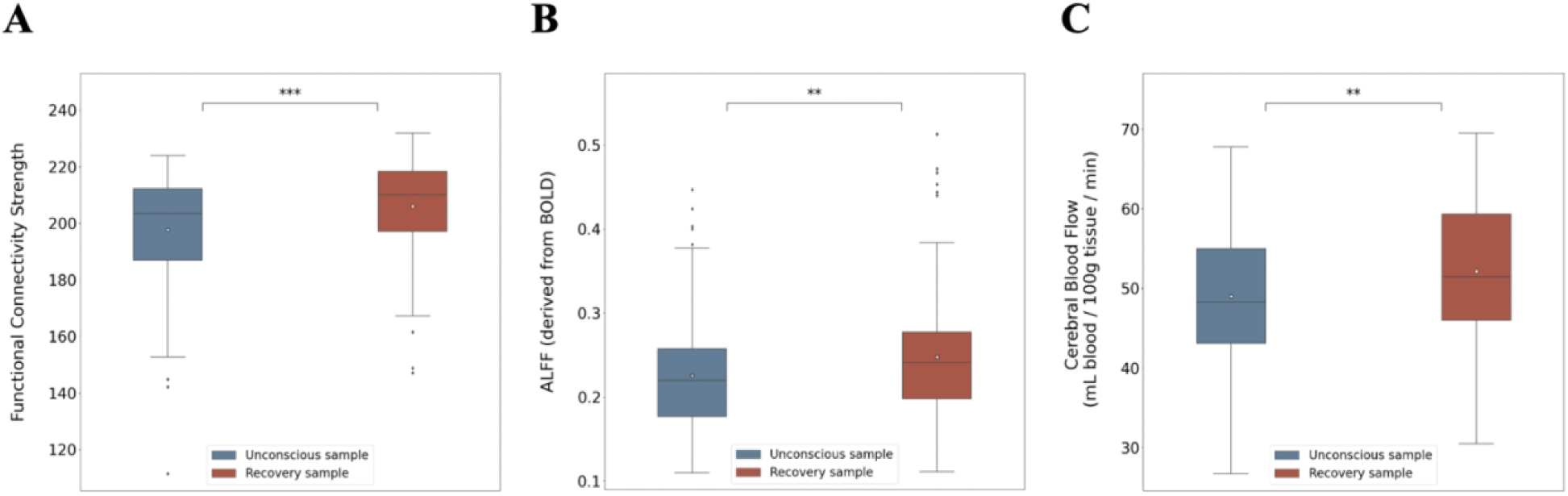
Whole-brain differences in connectivity and neurovascular variables between the subset of individuals who remained unconscious until the end of the experiment (unconscious sample) and the individuals who started regaining consciousness earlier (recovery sample). **A**: Differences in whole-brain functional connectivity strength between the two samples (n = 132 brain regions/datapoints for each boxplot). **B**: Differences in whole-brain ALFF between the two samples (n = 132 brain regions/datapoints for each boxplot). **C**: Differences in whole-brain cerebral blood flow between the two samples (n = 132 brain regions/datapoints for each boxplot). *p*-value annotation legend: non-significant (ns): 0.05 < *p* ≤ 1, *: 0.01 < *p* ≤ 0.05, **: 0.001< *p* ≤ 0.01, ***: 10^-4^ < *p* ≤ 10^-3^, ****: *p* ≤ 10^-4^; *p*-values here correspond to two-sided Mann-Whitney U tests.

Despite differences in functional connectivity strength between the two samples, there were no significant differences in SFC between the two samples, either at the whole-brain or at the network level.

In neurovascular dynamics, individuals who regained consciousness early had significantly higher levels of awake ALFF (percentage difference: 9.6% two-sided Mann-Whitney U test: *p* = 0.003; **Figure 7B**) and CBF (6.2% increase; *p* = 0.004; **Figure 7C**) across the whole brain, compared to the individuals who remained unconscious until the end of the experimental session. Despite these global differences across the two samples, there were no specific networks that displayed significant changes in neurovascular dynamics. Lastly, there were no significant differences in fractional ALFF between the two samples.

### Relationship between connectivity and neurovascular dynamics

In previous sections, we have established changes in connectivity and neurovascular dynamics taking place as the brain transitions between wakefulness, dexmedetomidine-induced unconsciousness, and the early-stage of consciousness recovery. Is there, however, a correspondence between these two biological aspects? Do changes in connectivity correlate with changes in neurovascular dynamics? To address this question, we focus on the relationship between functional connectivity and SFC with the two neurovascular markers of interest: ALFF and CBF.

First, we observed a significant negative correlation between a brain region’s functional connectivity strength and its average ALFF (permutation test across subjects: mean Spearman’s *r* across subjects = -0.19; *p_SPICE_* = 0.007) during the awake state. This result indicates that brain regions with higher overall ALFF tend to display lower overall functional connectivity with the rest of the brain, during wakefulness. This association did not persist during hypnosis.

In contrast to functional connectivity strength, a brain region’s SFC was negatively correlated with its average ALFF (permutation test across subjects: mean Spearman’s *r* across subjects = -0.12; *p_SPICE_* = 0.012), only during hypnosis. This result indicates that during hypnosis, brain regions with higher overall ALFF tend to display functional connectivity patterns that deviate more from the underlying structural connectivity patterns. Similarly, a brain region’s SFC was positively correlated with its average fractional ALFF (permutation test across subjects: mean Spearman’s *r* across subjects = 0.32; *p_SPICE_* = 0.029) during hypnosis.

Lastly, we observed that a brain region’s functional connectivity strength during wakefulness predicted the subsequent percentage change in CBF in that region, during the transition from wakefulness to hypnosis (permutation test across subjects: mean Spearman’s *r* across subjects = 0.08; *p_SPICE_* = 0.043).

## DISCUSSION

Characterizing the neurophysiological changes underlying loss and recovery of consciousness is a fundamental goal in neuroscience. In this work, we examine how the human brain alters its connectivity and neurovascular patterns across these transition periods, by analyzing diffusion-weighted, resting-state functional MRI, and ASL imaging data acquired at multiple time-points during dexmedetomidine-induced unconsciousness. Our results are seven-fold: (1) Functional connectivity strength prominently decreases across the whole brain from wakefulness to hypnosis; (2) SFC significantly increases from wakefulness to hypnosis, and starts decreasing as individuals start regaining consciousness, especially across cerebellar connectivity; (3) ALFF significantly increases across the brain during the wakefulness-hypnosis transition, and especially within visual and somatomotor regions; (4) CBF robustly decreases across the brain during the wakefulness-hypnosis transition—especially within the brainstem, thalamus, and cerebellum—and continues decreasing even while the brain starts regaining consciousness; (5) earlier recovery of consciousness is associated with higher baseline (awake) levels of functional connectivity strength, ALFF, and CBF, as well as female sex; (6) during wakefulness, brain regions with higher ALFF display lower overall functional connectivity with the rest of the brain; during hypnosis, brain regions with higher ALFF display weaker SFC; and lastly (7) brain regions with stronger levels of functional connectivity strength during wakefulness lower their CBF more as they enter hypnosis. In the paragraphs below, we discuss each finding.

### Connectivity changes

The transition from wakefulness to dexmedetomidine-induced unconsciousness, and then recovery of consciousness, was characterized by widespread changes in functional connectivity strength across the whole brain. During loss of behavioral responsiveness, functional connectivity strength—assessed via ASL imaging—dropped across all resting-state networks examined (visual, somatomotor, dorsal attention, ventral attention, fronto-parietal, default mode, limbic, subcortical, and cerebellar), indicating that networks entering hypnosis become less coherent with the rest of the brain across the frequency band assessed by resting-state fMRI. After hypnosis, whole-brain functional connectivity strength began increasing as the individuals started regaining consciousness, even before behavioral arousal. These findings are consistent with previous work utilizing a complementary definition of functional connectivity strength derived from resting-state BOLD fMRI alongside a set of functional networks (subcortical, sensory, default mode, attention/executive, language/memory), reporting a significant reduction followed by a non-significant increase in functional connectivity strength, during the wakefulness-to-unconsciousness and unconsciousness-to-recovery transitions, respectively.^14^ Building on this work, we additionally report significant increases in functional connectivity strength during the transition from hypnosis to recovery, primarily within the visual and default mode networks.

We next turned to changes in the coupling between functional connectivity and the underlying structural connectivity, as individuals traversed from wakefulness to hypnosis and finally to early-stage recovery. SFC robustly increased during loss of consciousness, especially across the unimodal sensory and motor, higher-order transmodal (fronto-parietal and default mode), and cerebellar networks. As individuals began recovering consciousness, whole-brain SFC started decreasing towards awake levels; such decreases were predominantly robust across the cerebellum (for a more expansive discussion on the putative role of the cerebellum in conscious-unconscious transitions, see the corresponding section in our **Supplementary Material**).

Intuitively, a brain region’s SFC captures the extent to which its functional connectivity patterns mirror the underlying structural connectivity patterns.^43^ Thus, the observed increase in whole-brain SFC from wakefulness to hypnosis may reflect a decrease in the number of functional configurations the brain can attain during loss of consciousness, given the underlying anatomical connectivity. Our results complement prior work identifying a smaller number of functional configurations and functional connectivity patterns that were progressively more similar to the underlying structural connectivity in patients with disorders of consciousness (vegetative state/unresponsive wakefulness syndrome or in minimally conscious state),^44–46^ in macaque monkeys anesthetized with propofol,^47,48^ sevoflurane,^48^ or ketamine,^48^ in humans transitioning from the awake to propofol-induced unconscious state,^46,49^ and in humans transitioning from wakefulness to deeper levels of sleep.^49,50^

### Neurovascular changes

We also assessed changes in neurovascular dynamics, as individuals transitioned from wakefulness to dexmedetomidine-induced unconsciousness, and finally early-stage recovery. To that end, we focused on two markers: CBF (derived from ASL imaging) and ALFF (derived from resting-state fMRI). ASL-derived CBF provides a direct measure of brain perfusion, which reflects a combination of cerebrovascular integrity, cerebral autoregulation, and neurovascular coupling.^51,52^ In our cohort of young, healthy subjects in whom cerebrovascular disease is highly unlikely, changes in CBF should predominantly reflect neural activity. ALFF provides a measure of neurovascular dynamics as assessed by fluctuations in the BOLD signal, which is sensitive to changes in CBF, cerebral blood volume, and cerebral oxygen metabolism.^53^ Although CBF and ALFF have been shown to be correlated, they may also become decoupled.^16^

We observed significant decreases in CBF, during dexmedetomidine-induced unconsciousness, across all networks and almost all brain regions examined. Regions where CBF decreased in magnitude included the brainstem, thalamus, and cerebellum. Interestingly, CBF continued decreasing across all brain networks even during the early stages of recovery of consciousness; this is in stark contrast to the connectivity markers (functional connectivity strength and SFC) which started reverting back to their awake levels, even before behavioral arousal. This finding could point towards changes in CBF as not being sufficient conditions for the recovery of consciousness.

Our results recapitulate and expand upon complementary studies identifying widespread decreases in cortical and subcortical CBF across a variety of different dosages of dexmedetomidine administration.^54,55^ Furthermore, our findings strongly align with the results of a previous study which not only reported significant decreases in CBF across the thalamus and higher-order transmodal networks (i.e., default mode and fronto-parietal) during dexmedetomidine-induced unconsciousness, but also revealed that these decreases in CBF did not immediately revert to their awake levels during recovery of consciousness.^12^

To better understand why dexmedetomidine reduces CBF, we turn to its role as an α_2_-adrenergic receptor agonist. Agonists, such as dexmedetomidine, have been reported to decrease CBF by (i) activating centrally and peripherally located α_2A_-adrenergic receptors mediating sympatholytic and vagomimetic effects, causing a decrease in heart rate and arterial pressure,^56,57^ and/or (ii) activating α_2B_-adrenergic receptors in vascular smooth muscle, causing cerebral vasoconstriction.^54,56^ In addition to decreasing during the transition from wakefulness to hypnosis, our analyses indicated that CBF continued decreasing in magnitude even during the initial stages of recovery of consciousness. This finding is physiologically supported by work demonstrating that dexmedetomidine weakens cerebral autoregulation—the ability of cerebral vasculature to maintain stable CBF despite changes in arterial pressure—causing a delay in CBF restoration back to awake levels during conditions of already reduced CBF (such as during dexmedetomidine-induced unconsciousness).^58^ Besides a potentially weakened cerebral autoregulation, sustained reductions in CBF might also reflect altered neurovascular coupling due to the reduced noradrenergic tone. By decreasing norepinephrine release in arousal-promoting regions, dexmedetomidine could be momentarily weakening the temporal coupling between oxygen demand and blood supply across these regions,^59^ thus delaying the return of CBF to awake levels.

We next turn to the second neurovascular marker of interest, ALFF. Much like CBF,^60–63^ ALFF is a metric with high test-retest reliability.^64,65^ ALFF captures the local power of low-frequency fluctuations within the 0.01–0.08 Hz range of the resting-state BOLD signal; these fluctuations are thought to reflect each brain region’s intrinsic, spontaneous neural activity, relatively devoid of cognitive load.^42,66^ In contrast to CBF, ALFF robustly increased across the whole brain as individuals transitioned from wakefulness to hypnosis. Fractional ALFF, on the other hand, which measures the relative power of low frequency fluctuations compared to the overall BOLD signal, decreased across the whole brain. On the regional level, ALFF increases (and similarly fractional ALFF decreases) were dominant within unimodal regions involved in visual, somatomotor, and auditory processing.

Our ALFF results align with previous studies reporting increases in the power of low-frequency BOLD oscillations within a comparable frequency range (0.03–0.10 Hz) across unimodal sensory and motor regions, during the transition from wakefulness to sleep.^30,67^ Similarly, two other studies reported a significant increase in spontaneous, low-frequency BOLD fluctuations (0.01–0.08 Hz) from resting wakefulness to light sleep, especially within primary sensory areas including visual, motor, and auditory cortices, with the visual cortex displaying the most robust increases.^68,69^ In fact, the magnitude of low-frequency BOLD signal fluctuations observed in the visual cortex during sleep was comparable to that observed during the performance of a visual task while awake.^69^ The ALFF increases observed during both dexmedetomidine-induced unconsciousness and sleep may reflect changes in neural oscillations—and specifically increases in slow wave amplitudes—preceding and driving the clearance of metabolic waste from the brain—a process more commonly referred to as glymphatic clearance^70^—and could provide further evidence of dexmedetomidine acting as a potential glymphatic clearance enhancer^71–74^ in humans.

Interestingly, another study examining changes in ALFF using a different anesthetic—propofol— reported overall decreases in both ALFF and fractional ALFF during light and deep sedation, compared to wakefulness, and especially within transmodal frontal cortices.^75^ Collectively, these findings could point towards dexmedetomidine and natural sleep imprinting similar effects on the cortical BOLD signal during the unconscious state—especially within unimodal sensory cortices—as well as highlight the contrasting effects between dexmedetomidine/sleep and propofol in inducing unconsciousness, with the latter preferentially altering the BOLD signal of transmodal cortices.

Collectively, the widespread increases in ALFF, decreases in fractional ALFF, and decreases in CBF observed during dexmedetomidine-induced hypnosis could support the notion that the brain enters a metabolically less demanding state. Arriving at this conclusion requires revisiting the physiological basis of the BOLD signal. The BOLD signal obtained from resting-state functional MRI is sensitive to local changes in the concentration of deoxygenated hemoglobin in the blood, which acts as a paramagnetic contrast agent causing inhomogeneities in the magnetic field and decreasing MRI signal intensity.^76–78^ Given that the BOLD signal fundamentally captures changes in brain blood deoxyhemoglobin, it has been used to indirectly infer changes in local neuronal activation, via neurovascular coupling: when neurons in a brain region are activated, oxygen is extracted from the local vasculature, leading to an increase in the concentration of deoxygenated hemoglobin. Therefore, the balance between oxygen consumption and oxygen supply modulates the BOLD signal. Prior work in humans has demonstrated that this balance shifts during both NREM sleep^79^ and after administration of dexmedetomidine,^55^ with oxygen consumption decreasing proportionately more than oxygen supply, compared to wakefulness. This leads to a decrease in paramagnetic deoxyhemoglobin, and, in turn, an increase in the BOLD signal. The increased ALFF and decreased CBF observed in our work could thus similarly suggest a decrease in oxygen supply accompanied by an even larger decrease in oxygen consumption taking place during dexmedetomidine-induced hypnosis, consistent with the brain attaining a metabolically less demanding state. This notion is also corroborated by the decreased fractional ALFF we observed during hypnosis, considering recent work reporting strong correlations between fractional ALFF with oxygen and glucose metabolism.^80^

### Individuals who remained unconscious versus individuals who regained consciousness

Given that a subset of the individuals in our sample unexpectedly regained consciousness before the end of the experimental session while still receiving a maintenance infusion of dexmedetomidine dosed by weight (see **Methodological Limitations** section in our **Supplementary Material**), we attempted to identify whether any of the aforementioned connectivity and neurovascular markers differed between them (recovery sample) and the individuals who remained unconscious (unconscious sample).

Indeed, the recovery sample displayed significantly higher baseline (awake) levels of mean functional connectivity strength compared to the rest of the individuals, especially within the visual, default mode, and cerebellar networks. The same sample also displayed higher levels of awake ALFF and CBF across the whole brain, compared to the unconscious sample. These differences could intuitively shed light on why the former group might have been able to wake up earlier: a decrease in these individuals’ functional connectivity strength and CBF during hypnosis could still provide them with more integrative capacity and neurovascular tone compared to the rest of the individuals, due to their higher baseline values. Interestingly, all individuals within the recovery sample were females, an observation in accordance with recent work in the field indicating that females regain consciousness and recover cognition faster than males after anesthesia—a process appearing to be modulated by testosterone.^81–83^ More generally, dexmedetomidine has been shown to differentially impact females versus males, producing, for instance, a weaker morphine-sparing effect in controlling post-operative acute pain in females.^84^ Critically, investigating the sex differences between commonly used anesthetic agents—and how their differential impact depends on other factors known to alter induction and emergence timescales, such as age and health status—would be a particularly fruitful clinical direction.

### Relationship between connectivity and neurovascular dynamics

Finally, we examined whether the overall changes in connectivity observed during the transition from wakefulness to dexmedetomidine-induced unconsciousness, and finally early-stage recovery, correlated with the reported changes in neurovascular dynamics. Indeed, brain regions characterized by higher levels of spontaneous BOLD fluctuations *during wakefulness*—in the form of higher awake ALFF—tended to be less functionally connected to the rest of the brain. Instead, brain regions with higher levels of spontaneous BOLD fluctuations *during hypnosis* displayed lower SFC: that is, functional connectivity patterns that deviated more from the underlying anatomical connectivity. These results indicate that at each (un)conscious state, brain regions with putatively higher metabolic needs become more functionally flexible and independent from other brain regions as well as from the underlying anatomical constrains. This could perhaps represent a neuroprotective mechanism to compensate for high awake metabolism, as such metabolically active regions would be highly susceptible in the presence of neurodegenerative disorders, such as Alzheimer’s disease, given their capacity to more effectively accelerate pathology.^85,86^

Lastly, baseline (awake) levels of functional connectivity strength predicted the percentage change in CBF during the transition from wakefulness to hypnosis. In other words, brain regions that were more strongly functionally connected to the rest of the brain tended to decrease their CBF more as they entered hypnosis. This finding could indicate once again how energetically consuming and topologically central brain regions might be entering a “resting” phase during unconsciousness, as a means to recalibrate and restore synaptic homeostasis.^87^

In summary, we analyzed magnetic resonance imaging data collected across multiple time-points to characterize how the human brain’s connectivity and neurovascular dynamics change as it transitions from wakefulness to dexmedetomidine-induced unconsciousness, and early-stage recovery of consciousness. During hypnosis, brain regions became less functionally synchronized to each other; they attained a smaller number of functional configurations compared to wakefulness, and displayed functional connectivity patterns that were more similar to the underlying structural connectivity. Furthermore, cerebral blood flow significantly decreased across the whole brain, and less metabolically demanding low frequency fluctuations in the hemodynamic signal became more prominent. Collectively, loss of consciousness was accompanied by widespread connectivity and neurovascular changes in the brain, characteristic of less metabolically demanding dynamics.

## MATERIALS AND METHODS

### DATASET

Healthy individuals (n = 14; mean age: 28.2 ± 5.6 years; age range: 22 – 39 years; 10 female) were recruited from the local community and scanned at the Hospital of the University of Pennsylvania using a Siemens Prisma 3T scanner (64-channel head/neck coil). They specifically underwent a battery of magnetic resonance imaging (MRI) sequences, including T1-weighted (voxel size: 0.8 mm isotropic; repetition time [TR]: 2400 ms; echo time [TE]: 2.24 ms), diffusion-weighted (diffusion imaging; voxel size: 3 mm isotropic; TR: 2500 ms; TE: 65 ms; 61 non-colinear directions; 8 b=0 acquisitions), resting-state functional (700 volumes; voxel size: 2 mm isotropic; TR: 720 ms; TE: 37 ms), and arterial spin labeling (single-shot background-suppressed pseudo-continuous arterial spin labeling [ASL]: 60 volumes; voxel size: 3.75 mm isotropic; TR: 4000 ms; TE: 10.03 ms, along with a reference image to estimate the equilibrium magnetization of blood [M0]) imaging. Informed consent was obtained from all individuals, and the procedures were approved by the University of Pennsylvania Institutional Review Board.

The experimental protocol followed is schematically illustrated in **Figure 1** and described below. Baseline scans (one T1-weighted scan, one resting-state functional MRI scan, and three pairs of diffusion-weighted scans each followed by an ASL sequence) were first acquired while each subject was awake. Each subject then received administration of the anesthetic agent dexmedetomidine (1μg/kg) via an intravenous catheter over 10 minutes followed by a dexmedetomidine infusion (1μg/kg/hr) for the duration of the remaining 3 scans to pharmacologically enter a state resembling NREM sleep. After the dexmedetomidine loading dose and during the dexmedetomidine infusion—while the subjects were unconscious—the same MRI sequences (with the exception of the T1-weighted sequence) were re-acquired.

Out of the total 14 individuals who completed the experiment described above, 8 (mean age: 25.6 ± 4.6 years; age range: 22–36 years; all female) began prematurely waking up or displaying signs of behavioral arousal (i.e., squeezing the alarm ball, opening eyes, moving extremities) during the ongoing dexmedetomidine infusion towards the end of the experiment, starting at the beginning of their 3^rd^ “hypnotic” diffusion-weighted scan (**Figure 1**). To further understand this small sub-sample of individuals who recovered consciousness early, we used their data in the “3^rd^ hypnotic” time-point to define an “early-stage recovery” time-point.

To summarize, data from all 14 individuals were used for the first 5 time-points (the 3 awake and first 2 hypnotic time-points), data from only the 6 individuals who remained unconscious until the end of the experiment were used to define a 3^rd^ hypnotic time-point, and data from only the 8 individuals who started regaining consciousness early were used to define an early-stage recovery time-point. Comparisons between the values of interest across the awake, hypnotic, and recovery time-points are described in our Results section.

### PROCESSING PIPELINES

For all analyses considered, cortical, subcortical, and cerebellar brain areas were defined using a combination of the Harvard-Oxford atlas and the AAL atlas. This combined Harvard-Oxford-AAL atlas incorporates 91 cortical regions of interest (ROIs) defined from *FSL*’s^88^ Harvard-Oxford maximum likelihood cortical atlas (HarvardOxford-cort-maxprob-thr25-1mm.nii), 15 subcortical ROIs from the Harvard-Oxford maximum likelihood subcortical atlas (HarvardOxford-sub-maxprob-thr25-1mm.nii), and 26 cerebellar ROIs from the Automated Anatomical Labeling atlas (AAL), for a total of 132 brain regions.^89–93^

We applied two pipelines: a (1) <u>connectivity pipeline</u>, wherein changes in functional connectivity and the statistical correlation between structural and functional connectivity—more commonly referred to as structure-function coupling—were assessed; and a (2) <u>neurovascular dynamics</u> <u>pipeline</u>, wherein changes in two markers—ALFF and CBF—were estimated. We describe both pipelines in detail below:

## I. Connectivity pipeline

### Structural connectivity

Structural connectivity was assessed using the acquired diffusion-weighted imaging sequences. Diffusion data were processed using the *MRtrix3* toolbox.^94^ The first preprocessing step was denoising the data and manually examining the residual maps (difference maps between the denoised and original diffusion images) to ensure that no particular brain region was disproportionately impacted by artifactual noise. Eddy current-induced distortion and motion correction were then applied on the denoised diffusion data using the *dwifslpreproc* command. After this step had completed, the percentage of outlier slices for each diffusion scan was output and manually inspected; all diffusion scans used contained less than 2% of slices classified as outliers. Finally, B_1_ field inhomogeneity correction was applied on the diffusion data and an accurate brain mask was generated, to restrict all analyses just to brain voxels.

The second part of diffusion data processing included determining the orientation of diffusion within each brain voxel. First, a basis response function was generated from each individual’s diffusion scans, using the *dwi2response* command and the “dhollander” algorithm; the purpose of this step was to decompose the diffusion signal within each tissue type (gray matter, white matter, and CSF) into a set of fiber orientations across different directions. These basis response functions were subsequently used to estimate fiber orientation densities using multi-shell, multi-tissue constrained spherical deconvolution (command: *dwi2fod*). The resulting fiber orientation densities were then normalized to ensure that any observed differences were not due to intensity contrasts in the diffusion image.

The last component of the diffusion data processing invoked reconstructing the white matter tracts (i.e., streamlines) connecting the different gray matter brain regions, using anatomically constrained probabilistic tractography. For this purpose, an initial whole-brain tractogram containing ten million streamlines was first generated using the command *tckgen*; the “-act” flag was supplied to the command to only focus on streamlines constrained to the white matter. The streamlines were then refined by assigning them weights to reduce known biases (SIFT2 method;^95^ command *tcksift2*). Finally, each subject’s refined streamline map was registered to a parcellated brain atlas (Harvard-Oxford-AAL atlas: n = 132 brain regions) that had been registered to the subject’s native anatomical space. Subject-specific, symmetric, weighted structural connectivity matrices of size 132 x 132 were then generated for each diffusion scan; each row and column of the matrix corresponded to a brain region and each edge to the SIFT2-weighted sum of the streamlines connecting any two given regions divided by the sum of the regions’ gray matter volumes.

### Functional connectivity

Functional connectivity was assessed using the ASL sequences acquired, a previously validated methodology,^40,41^ during the 6 different experimental time-points. ASL data were processed using *MATLAB* (version R2021a: The MathWorks, Inc.) and *SPM* (version 12)^96^. The ASL images were first motion corrected using the commands *spm_realign* (additional parameter used: 5 mm full-width half-maximum Gaussian kernel smoothing applied) and *spm_reslice*. The motion corrected ASL images, as well as the reference image M0 acquired at each time-point, were then co-registered to the anatomical space (the individual’s T1-weighted image) using trilinear interpolation, and lastly to the standardized MNI (2 mm) space using non-linear registration (*FSL*’s *fnirt* command).

Using the processed ASL images, we finally constructed functional connectivity matrices for each of the 6 time-points considered. The mean ASL regional time-series (size: 1 x 60 volumes) was computed for each brain region of interest (for a total of 132 brain ROIs, as per the Harvard-Oxford-AAL atlas); the coherence between two brain regions’ ASL regional time-series was calculated.^97^ From a technical standpoint, coherence measures the amount of cross-correlation between two time-series in the frequency domain. Thus, the coherence between brain region *i* and brain region *j* at frequency *f* is defined as:

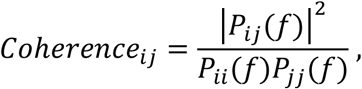

where *P*_*ij*_(*f*) is the cross power spectral density between the time-series corresponding to brain regions *i* and *j* at frequency *f*.^97,98^ The average coherence between two brain regions across all frequencies greater than 0.01 Hz was computed using *MATLAB*’s *mscohere* command, and was used to define the functional connectivity between these two brain regions. A lower frequency threshold of 0.01 Hz was used to remove signals of non-biological origin.^99^ After computing the coherence between all pairs of brain regions for each ASL scan acquired, we constructed the corresponding functional connectivity matrices. A Fisher r-to-z transform was lastly applied on the functional connectivity matrices, using *MATLAB*’s *atanh* function, to normalize their values.^98^

After the functional connectivity matrices were constructed for each individual, each brain region’s <u>functional connectivity strength</u> was calculated and defined as the sum of all edges between that brain region and all other brain regions in the functional connectivity matrix.

### Structure-function coupling

Both structural and functional connectivities were assessed during all awake and hypnotic time-points; given that there were three pairs of structural connectivity matrices generated at the awake state and three pairs at the hypnotic state, while there were only three functional connectivity matrices, we selected the three structural connectivity matrices at each state corresponding to the diffusion scans with the least amount of noise (as defined by each scan’s percentage of outlier slices, as defined above). To examine the statistical relationship between structural and functional connectivity across each of these time-points, we computed their structure-function coupling (SFC). A brain region’s structure-function coupling was defined as the Pearson’s correlation coefficient between its structural connectivity vector (size: 1 x 132 ROIs) and its functional connectivity vector (size: 1 x 132 ROIs) towards all other brain regions, after excluding the self-connection and any other entries where either the regional structural or functional connectivity was equal to zero, as previously defined.^43^

These processes allowed us to compute six types of metrics for each connectivity metric (functional connectivity strength and SFC) examined: (i) the metric corresponding to each of the 6 time-points, (ii) the metric during the average awake state, (iii) the metric during the average hypnotic state, (iv) the metric during the recovery state, (v) the percent change in the metric from the average awake to the average hypnotic state, and (vi) the percent change in the metric from the last hypnotic time-point to the recovery time-point. The latter two were calculated using the formulae:

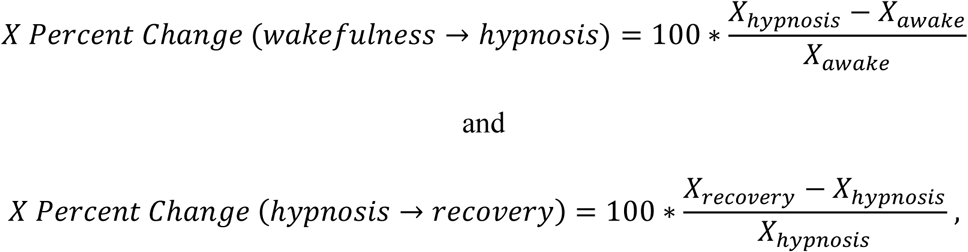

where *X* denotes the connectivity metric of interest (X = functional connectivity strength or SFC).

## II. Neurovascular dynamics pipeline

### Amplitude of low-frequency fluctuations (ALFF)

Changes in ALFF for each individual were assessed using the resting-state functional MRIs acquired during their awake and hypnotic states, at the beginning and end of the experimental setup, respectively (**Figure 1**). The resting-state functional MRIs were processed with *CONN* (https://web.conn-toolbox.org/home; version 20.b),^100,101^ using its “default pre-processing pipeline for volume-based analyses (direct normalization to MNI-space).” A more detailed description of the pipeline has been previously reported;^43^ here we provide a brief listing of the processing steps: (i) Each subject’s functional scans were co-registered to a reference image, (ii) corrected for any temporal misalignment between slices that may have occurred during acquisition, (iii) the structural (T1-weighted) scans were segmented into gray matter, white matter, and CSF tissue classes, and both structural and functional scans were normalized into standardized MNI space, (iv) the functional images were smoothed using an 8 mm full-width half-maximum Gaussian kernel to increase the BOLD signal-to-noise ratio.^101^ After these pre-processing steps had taken place, *CONN*’s default denoising pipeline was utilized to remove any potential confounders identified in the BOLD signal. Confounders included noise components from white matter and cerebrospinal areas, estimated subject motion parameters (i.e., 3 rotation and 3 translation parameters, and their 6 associated first-order derivatives), flagged outlier scans, and session-related effects such as constant and linear BOLD signal trends. Frequencies below 0.008 Hz were also removed to mitigate the presence of low-frequency drifts and signals of non-biological origin. B0 field inhomogeneities in each subject’s resting-state functional MRI were also corrected, using double-echo gradient echo sequences (2 magnitude volumes and 1 phase difference volume [voxel size: 3 mm isotropic; TR: 580 ms; TE difference: 2.46 ms]) acquired during the MRI battery.

After extracting each individual’s denoised resting-state BOLD signal time-series, the ALFF was estimated at each experimental time-point. The ALFF captures the power of low frequencies (0.01–0.08 Hz) within the BOLD signal, and was computed by (i) bandpass filtering the denoised BOLD signal time-series of each brain region over the frequency range 0.01–0.08 Hz, and then computing the standard deviation of that filtered BOLD signal.^101,102^ This allowed us to estimate an ALFF value for each brain region (as defined by the Harvard-Oxford-AAL atlas: n = 132 brain regions), for each individual, and at each experimental time-point.

Similarly, a brain region’s fractional ALFF was defined as the ratio between its ALFF and the standard deviation of the overall BOLD signal (frequencies ≥ 0.008 Hz). In other words, fractional ALFF quantifies the power of the BOLD signal within the low-frequency range, relative to the power of the overall BOLD signal.

There were six types of metrics extracted for each brain region and for each individual: (i) the (fractional) ALFF during the awake state, (ii) the (fractional) ALFF during the hypnotic state, and the percent change in (fractional) ALFF from the awake to the hypnotic state. The latter two were calculated using the formula:

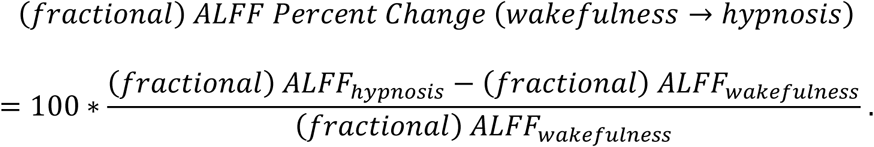

Awake levels of (fractional) ALFF were computed across all individuals (n = 14) and hypnotic levels of (fractional) ALFF were computed only across the individuals who remained unconscious until the end of the experimental session (n = 6; see the “Dataset” section). Given that there were only two resting-state functional MRI sequences that were acquired for each individual during the experiment—once during wakefulness and once at the end of the experimental setup (**Figure 1**)— we were not able to extract the percent change in (fractional) ALFF from a hypnotic time-point to a recovery time-point (see **Supplementary Material: Methodological Limitations**).

### Cerebral blood flow

CBF within each brain region was estimated using the processed ASL sequences (see ‘Functional Connectivity’ section above). A CBF map corresponding to each of the 6 experimental time-points was generated using the equation:

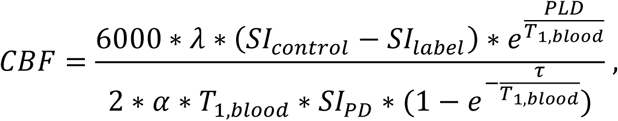

where *λ* is the brain/blood partition coefficient measured in mL/g (set to 0.9), *SI*_*control*_ and *SI*_*label*_ are the time-averaged signal intensities in the control and label images extracted from the processed ASL image (note: the ASL volumes consist of control and label images acquired in pairs and identically scaled), *PLD* refers to the post-labeling delay in the ASL scan measured in seconds (set to 1.5), *T*_1,*blood*_ is the longitudinal relaxation time of blood measured in seconds (set to 1.65), *α* is the labeling efficiency (scalar; set to 0.72), *SI*_*PD*_ is the time-averaged signal intensity of the proton density-weighted image (M0 reference image), and *τ* is the label duration measured in seconds (set to 1.5).^103^ The 6000 factor in the numerator is included to convert the units of the above equation from mL/g/seconds to the more conventionally used mL/100 g/min.^103^

After the above calculation was performed for each time-point, six types of metrics were extracted for each brain region and for each individual: (i) the CBF corresponding to each of the 6 time-points, (ii) the CBF during the average awake state, (iii) the CBF during the average hypnotic state, the CBF during the hypnotic state, (v) the percent change in CBF from the average awake to the average hypnotic state, and (vi) the percent change in CBF from the last hypnotic time-point to the recovery time-point. The latter two were calculated using the formulae:

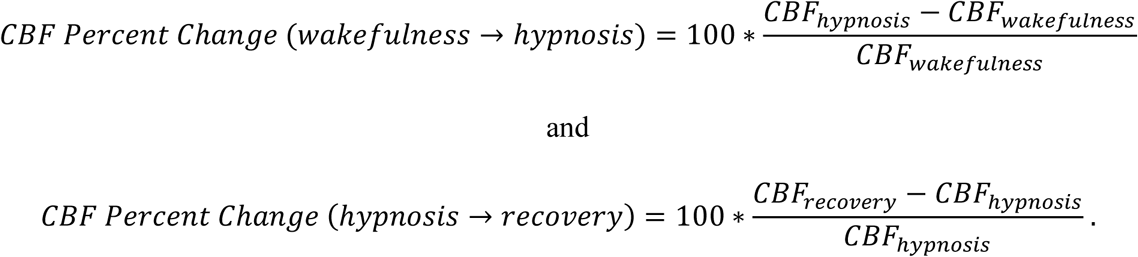

## STATISTICAL ANALYSES

Statistical analyses were performed using *MATLAB* (version R2021a: The MathWorks, Inc.) and *Python* (version 3.7). A *p*-value < 0.05 was used as the threshold for significance.

### Whole-brain level analyses

Repeated measures one-way analysis of variance (rmANOVA) tests were used to statistically compare the mean values of functional connectivity strength, SFC, and CBF across the 5 experimental time-points (3 awake time-points and the first 2 hypnotic time-points) described in the “Whole-brain level” sections of our Results. The rmANOVA tests were followed by *post-hoc* correction for multiple comparisons (Tukey’s honestly significant difference [HSD] test) to identify which pairs of time-points significantly differed in their mean values (as shown in **Figures 2A, 2C, 5A**). To compare the mean values of the metrics of interest between the overall awake state (i.e., the average metric value across the 3 “awake” time-points) and the overall hypnotic state (i.e., the average metric value across the first 2 “hypnotic” time-points) (as shown in **Figures 2B, 2D, 4A, 4B**), we used a Wilcoxon test for paired measurements, a non-parametric paired test bearing no assumption on the normality of the data distribution. We used the same test to compare brain regions’ functional connectivity strength, SFC, and CBF between the 2^nd^ hypnotic time-point and recovery time-point (as shown in Figure **5B**). Data from the 3^rd^ hypnotic time-point were not included in the rmANOVA analyses, as a subset of the individuals (n = 8) did not remain unconscious during that time-point.

Lastly, paired *t*-tests were used to compare the global functional connectivity strength, SFC, ALFF, and CBF values between each individual’s average awake vs. average hypnotic, and last hypnotic time-point vs. recovery time-point (as shown in **Figures 3, 4C, 4D, 5C**). For each comparison, we report the percent change in the variables’ magnitude and the corresponding *p*-value.

### Network level analyses

To compare the mean functional connectivity strength, SFC, ALFF, and CBF metrics between the awake, hypnotic, and recovery states (as shown in **Supplementary Figures 1-5, 7-10**) across the 9 networks examined (visual, somatomotor, dorsal attention, ventral attention, limbic, fronto-parietal, default mode, subcortical, and cerebellar networks)^37^, we once again used paired samples Wilcoxon tests—with an additional Benjamini-Hochberg correction for multiple comparisons to control for the false discovery rate across these 9 comparisons. For each comparison, we report the percent change in the variables’ magnitude between the three states across each network and the corresponding *p*-value.

### Regional analyses

Paired *t*-tests were once again used to compare regional differences in functional connectivity strength, SFC, ALFF, and CBF between the awake, hypnotic, and recovery states. For each comparison, we report the percent change in the variables’ magnitude between the three states and the corresponding *p*-value.

### Individuals who remained unconscious versus individuals who regained consciousness

To compare the mean values of functional connectivity strength, SFC, and CBF between the subset of individuals who started regaining consciousness before the end of the experiment (recovery sample) and the rest of the individuals who remained unconscious (unconscious sample), we used a two-sided Mann-Whitney U test, a non-parametric test bearing no assumption on the normality of the data distribution (as shown in **Figure 7**). The same test—with Benjamini-Hochberg correction for multiple comparisons to control for the false discovery rate—was used to compare the two samples across the 9 networks examined (as shown in **Supplementary Figure 10**). In contrast to the previous three sections in our statistical analyses where we report percent changes in the variables’ magnitude, here we report the percent difference between the variables’ magnitude, since the two samples compared do not consist of the same individuals. The formula used to compute the percent difference for variable *X* (where *X* can be functional connectivity strength, SFC, ALFF, or CBF) was:

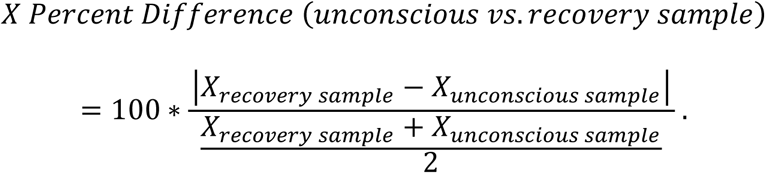

### Relationship between connectivity and neurovascular dynamics

The comparisons reported in the “Relationship between connectivity and neurovascular dynamics” section of our Results were carried out using a previously established “simple permutation-based test of intermodal correspondence” (SPICE test) that assesses how two brain maps correlate at the individual level, on average.^104^ We briefly describe the SPICE test below.

Let us assume that we want to correlate variables (i.e., brain maps) **X** and **Y** across multiple individuals. First, we format **X** and **Y** to each be of size <number of individuals x number of brain regions considered>, which in our case was 14 x 132. Each entry in these matrices corresponds to the value of the respective variable for a given individual and a given brain region. The permutation test then performs the following steps: (1) First, it computes the Spearman’s correlation coefficient between entries of each row of **X** and those in each row of **Y** (i.e., brain regions for each individual); (2) the average correlation coefficient across all individuals is then estimated, and designated as the observed correlation coefficient *C_obs_*; (3) the rows of **Y** are randomly sampled without replacement, generating a shuffled version of variable **Y** denoted: **Y_permuted_**; what this step does is shuffle the order of individuals within variable **Y**, while maintaining the order of individuals in variable **X** (original order of rows of **X**). This step also maintains inherent spatial dependencies within both **X** and **Y** (i.e., original order of columns of both **X** and **Y**); (4) the new average correlation coefficient across individuals between variables **X** and **Y_permuted_** is re-estimated as per steps 1 and 2, and designated as a null correlation coefficient *C_null_*; (5) steps 3 and 4 are repeated K times (here, K = 10,000) to generate a distribution of null coefficients; (6) a *p*-value (*p_SPICE_*) is finally estimated by counting the number of permutations where the (absolute) magnitude of *C_null_* exceeds that of *C_obs_*, divided by the total number of permutations (here 10,000) plus 1, which accounts for the identity permutation.^104^

In the statistical comparisons mentioned in the section “Relationship between connectivity and neurovascular dynamics” of our Results, we report (i) the average Spearman’s correlation coefficient *r* across subjects for each comparison (variable *C_obs_* described above) and (ii) the corresponding *p_SPICE_* value.

## CITATION DIVERSITY STATEMENT

Recent work in several fields of science has identified a bias in citation practices such that papers from women and other minority scholars are under-cited relative to the number of such papers in the field.^105–109^ We obtained the predicted gender of the first and last author of each reference by using databases that store the probability of a first name being carried by a woman.^109^ By this measure (and excluding self-citations to the first and last authors of our current paper), our references contain 3% woman(first)/woman(last), 12.9% man/woman, 28.7% woman/man, and 55.4% man/man. This method is limited in that a) names, pronouns, and social media profiles used to construct the databases may not, in every case, be indicative of gender identity and b) it cannot account for intersex, non-binary, or transgender people. We look forward to future work that could help us better understand how to support equitable practices in science.

## ACKNOWLEDGEMENTS

This work was supported by NIH grants T32-EB020087 (P.F.), R35GM142712 (A.R.M.-W.), R37MH125829 (T.D.S.), R01MH113550 (T.D.S. & D.S.B.), RF1MH116920 (T.D.S. & D.S.B.), R21MH106799 (T.D.S. & D.S.B.), R01MH112847 (R.T.S. & T.D.S.), R01MH123550 (R.T.S.), R01NS112274 (R.T.S.), R01EB031080 (J.A.D.), the Swartz Foundation (D.S.B.), and the John D. and Catherine T. MacArthur Foundation (D.S.B.).

## REFERENCES

1. Hudetz, A. G. General Anesthesia and Human Brain Connectivity. Brain Connect. 2, 291–302 (2012).

2. Långsjö, J. W. et al. Returning from Oblivion: Imaging the Neural Core of Consciousness. J. Neurosci. 32, 4935–4943 (2012).

3. Shehabi, Y. et al. Early Sedation with Dexmedetomidine in Critically Ill Patients. N. Engl. J. Med. 380, 2506–2517 (2019).

4. Kaur, M. & Singh, P. M. Current role of dexmedetomidine in clinical anesthesia and intensive care. Anesth. Essays Res. 5, 128–133 (2011).

5. Huupponen, E. et al. Electroencephalogram spindle activity during dexmedetomidine sedation and physiological sleep. Acta Anaesthesiol. Scand. 52, 289–294 (2008).

6. Akeju, O. et al. Dexmedetomidine promotes biomimetic non-rapid eye movement stage 3 sleep in humans: A pilot study. Clin. Neurophysiol. Off. J. Int. Fed. Clin. Neurophysiol. 129, 69–78 (2018).

7. Chamadia, S. et al. Oral Dexmedetomidine Promotes Non-rapid Eye Movement Stage 2 Sleep in Humans. Anesthesiology 133, 1234–1243 (2020).

8. Scott-Warren, V. L. & Sebastian, J. Dexmedetomidine: its use in intensive care medicine and anaesthesia. BJA Educ. 16, 242–246 (2016).

9. Brown, E. N., Purdon, P. L. & Van Dort, C. J. General Anesthesia and Altered States of Arousal: A Systems Neuroscience Analysis. Annu. Rev. Neurosci. 34, 601–628 (2011).

10. Zhang, Z. et al. Neuronal ensembles sufficient for recovery sleep and the sedative actions of α2 adrenergic agonists. Nat. Neurosci. 18, 553–561 (2015).

11. Gilsbach, R. et al. Genetic Dissection of α2-Adrenoceptor Functions in Adrenergic versus Nonadrenergic Cells. Mol. Pharmacol. 75, 1160–1170 (2009).

12. Akeju, O. et al. Disruption of thalamic functional connectivity is a neural correlate of dexmedetomidine-induced unconsciousness. eLife 3, e04499 (2014).

13. Nelson, L. E. et al. The α2-Adrenoceptor Agonist Dexmedetomidine Converges on an Endogenous Sleep-promoting Pathway to Exert Its Sedative Effects. Anesthesiology 98, 428– 436 (2003).

14. Hashmi, J. A. et al. Dexmedetomidine disrupts the local and global efficiency of large-scale brain networks. Anesthesiology 126, 419–430 (2017).

15. Guldenmund, P. et al. Brain functional connectivity differentiates dexmedetomidine from propofol and natural sleep. Br. J. Anaesth. 119, 674–684 (2017).

16. Baller, E. B. et al. Developmental coupling of cerebral blood flow and fMRI fluctuations in youth. Cell Rep. 38, 110576 (2022).

17. Jin, M. et al. Disturbed neurovascular coupling in hemodialysis patients. PeerJ 8, e8989 (2020).

18. Chiacchiaretta, P., Cerritelli, F., Bubbico, G., Perrucci, M. G. & Ferretti, A. Reduced Dynamic Coupling Between Spontaneous BOLD-CBF Fluctuations in Older Adults: A Dual-Echo pCASL Study. Front. Aging Neurosci. 10, 115 (2018).

19. Archila-Meléndez, M. E., Sorg, C. & Preibisch, C. Modeling the impact of neurovascular coupling impairments on BOLD-based functional connectivity at rest. NeuroImage 218, 116871 (2020).

20. Akeju, O. et al. Spatiotemporal Dynamics of Dexmedetomidine-Induced Electroencephalogram Oscillations. PLoS ONE 11, e0163431 (2016).

21. Akeju, O. et al. A Comparison of Propofol- and Dexmedetomidine-induced Electroencephalogram Dynamics Using Spectral and Coherence Analysis. Anesthesiology 121, 978–989 (2014).

22. Lee, U. et al. Disruption of Frontal-Parietal Communication by Ketamine, Propofol, and Sevoflurane. Anesthesiology 118, 1264–1275 (2013).

23. Lewis, L. D., et al. Rapid fragmentation of neuronal networks at the onset of propofol-induced unconsciousness. Proc. Natl. Acad. Sci. 109, E3377–E3386 (2012).

24. Murphy, M. et al. Propofol anesthesia and sleep: a high-density EEG study. Sleep 34, 283–291A (2011).

25. Purdon, P. L., Sampson, A., Pavone, K. J. & Brown, E. N. Clinical Electroencephalography for Anesthesiologists: Part I: Background and Basic Signatures. Anesthesiology 123, 937–960 (2015).

26. Akeju, O. et al. Electroencephalogram Signatures of Ketamine-Induced Unconsciousness. Clin. Neurophysiol. Off. J. Int. Fed. Clin. Neurophysiol. 127, 2414–2422 (2016).

27. Li, D. & Mashour, G. A. Cortical dynamics during psychedelic and anesthetized states induced by ketamine. NeuroImage 196, 32–40 (2019).

28. Brown, E. N., Pavone, K. J. & Naranjo, M. Multimodal General Anesthesia: Theory and Practice. Anesth. Analg. 127, 1246–1258 (2018).

29. Warnaby, C. E., Sleigh, J. W., Hight, D., Jbabdi, S. & Tracey, I. Investigation of Slow-wave Activity Saturation during Surgical Anesthesia Reveals a Signature of Neural Inertia in Humans. Anesthesiology 127, 645–657 (2017).

30. Fultz, N. E. et al. Coupled electrophysiological, hemodynamic, and cerebrospinal fluid oscillations in human sleep. Science 366, 628–631 (2019).

31. Schiff, N. D., Nauvel, T. & Victor, J. Large-Scale Brain Dynamics in Disorders of Consciousness. Curr. Opin. Neurobiol. 0, 7–14 (2014).

32. Young, G. B. The EEG in Coma. J. Clin. Neurophysiol. 17, 473 (2000).

33. Pal, D. et al. Level of Consciousness Is Dissociable from Electroencephalographic Measures of Cortical Connectivity, Slow Oscillations, and Complexity. J. Neurosci. 40, 605–618 (2020).

34. Pal, D. et al. Differential Role of Prefrontal and Parietal Cortices in Controlling Level of Consciousness. Curr. Biol. 28, 2145–2152.e5 (2018).

35. Davis, C. J., Clinton, J. M., Jewett, K. A., Zielinski, M. R. & Krueger, J. M. Delta Wave Power: An Independent Sleep Phenotype or Epiphenomenon? J. Clin. Sleep Med. JCSM Off. Publ. Am. Acad. Sleep Med. 7, S16–S18 (2011).

36. Fotiadis, P. et al. Structure–function coupling in macroscale human brain networks. Nat. Rev. Neurosci. 1–17 (2024) doi:10.1038/s41583-024-00846-6.

37. Yeo, B. T. T. et al. The organization of the human cerebral cortex estimated by intrinsic functional connectivity. J. Neurophysiol. 106, 1125–1165 (2011).

38. Margulies, D. S. et al. Situating the default-mode network along a principal gradient of macroscale cortical organization. Proc. Natl. Acad. Sci. 113, 12574–12579 (2016).

39. Sydnor, V. J. et al. Neurodevelopment of the association cortices: Patterns, mechanisms, and implications for psychopathology. Neuron 109, 2820–2846 (2021).

40. Li, Z. et al. Effects of resting state condition on reliability, trait specificity, and network connectivity of brain function measured with arterial spin labeled perfusion MRI. NeuroImage 173, 165–175 (2018).

41. Dai, W., Varma, G., Scheidegger, R. & Alsop, D. C. Quantifying fluctuations of resting state networks using arterial spin labeling perfusion MRI. J. Cereb. Blood Flow Metab. 36, 463– 473 (2016).

42. Zou, Q.-H. et al. An improved approach to detection of amplitude of low-frequency fluctuation (ALFF) for resting-state fMRI: Fractional ALFF. J. Neurosci. Methods 172, 137– 141 (2008).

43. Fotiadis, P. et al. Myelination and excitation-inhibition balance synergistically shape structure-function coupling across the human cortex. Nat. Commun. 14, 6115 (2023).

44. Demertzi, A. et al. Human consciousness is supported by dynamic complex patterns of brain signal coordination. Sci. Adv. 5, eaat7603 (2019).

45. Perl, Y. S. et al. Low-dimensional organization of global brain states of reduced consciousness. Cell Rep. 42, 112491 (2023).

46. Luppi, A. I. et al. Distributed harmonic patterns of structure-function dependence orchestrate human consciousness. *Commun*. Biol. 6, 1–19 (2023).

47. Barttfeld, P. et al. Signature of consciousness in the dynamics of resting-state brain activity. Proc. Natl. Acad. Sci. 112, 887–892 (2015).

48. Luppi, A. I. et al. Local orchestration of distributed functional patterns supporting loss and restoration of consciousness in the primate brain. Nat. Commun. 15, 2171 (2024).

49. Castro, P. et al. Dynamical structure-function correlations provide robust and generalizable signatures of consciousness in humans. *Commun*. Biol. 7, 1–12 (2024).

50. Tagliazucchi, E., Crossley, N., Bullmore, E. T. & Laufs, H. Deep sleep divides the cortex into opposite modes of anatomical–functional coupling. Brain Struct. Funct. 221, 4221–4234 (2016).

51. Dolui, S. et al. Reliability of arterial spin labeling derived cerebral blood flow in periventricular white matter. Neuroimage Rep. 1, 100063 (2021).

52. Detre, J. A. et al. Noninvasive MRI evaluation of cerebral blood flow in cerebrovascular disease. Neurology 50, 633–641 (1998).

53. Simon, A. B. & Buxton, R. B. Understanding the dynamic relationship between cerebral blood flow and the BOLD signal: Implications for quantitative functional MRI. NeuroImage 116, 158–167 (2015).

54. Prielipp, R. C. et al. Dexmedetomidine-Induced Sedation in Volunteers Decreases Regional and Global Cerebral Blood Flow. Anesth. Analg. 95, 1052 (2002).

55. Drummond, J. C. et al. Effect of dexmedetomidine on cerebral blood flow velocity, cerebral metabolic rate, and carbon dioxide response in normal humans. Anesthesiology 108, 225– 232 (2008).

56. Talke, P. & Anderson, B. J. Pharmacokinetics and pharmacodynamics of dexmedetomidine-induced vasoconstriction in healthy volunteers. Br. J. Clin. Pharmacol. 84, 1364–1372 (2018).

57. Mikkelsen, M. L. G. et al. The effect of dexmedetomidine on cerebral perfusion and oxygenation in healthy piglets with normal and lowered blood pressure anaesthetized with propofol-remifentanil total intravenous anaesthesia. Acta Vet. Scand. 59, 27 (2017).

58. Ogawa, Y. et al. Dexmedetomidine Weakens Dynamic Cerebral Autoregulation as Assessed by Transfer Function Analysis and the Thigh Cuff Method. Anesthesiology 109, 642–650 (2008).

59. Bekar, L. K., Wei, H. S. & Nedergaard, M. The locus coeruleus-norepinephrine network optimizes coupling of cerebral blood volume with oxygen demand. J. Cereb. Blood Flow Metab. 32, 2135–2145 (2012).

60. Yang, F. N. et al. Test-retest reliability of cerebral blood flow for assessing brain function at rest and during a vigilance task. NeuroImage 193, 157–166 (2019).

61. Almeida, J. R. C. et al. Test-retest reliability of cerebral blood flow in healthy individuals using arterial spin labeling: Findings from the EMBARC study. Magn. Reson. Imaging 45, 26– 33 (2018).

62. Binnie, L. R. et al. Test–retest reliability of arterial spin labelling for cerebral blood flow in older adults with small vessel disease. Transl. Stroke Res. 13, 583–594 (2022).

63. Kyrou, A. et al. Test-retest reliability of resting-state cerebral blood flow quantification using pulsed Arterial Spin Labeling (PASL) over 3 weeks vs 8 weeks in healthy controls. Psychiatry Res. Neuroimaging 341, 111823 (2024).

64. Küblböck, M. et al. Stability of low-frequency fluctuation amplitudes in prolonged resting-state fMRI. NeuroImage 103, 249–257 (2014).

65. Vedaei, F., Alizadeh, M., Romo, V., Mohamed, F. B. & Wu, C. The effect of general anesthesia on the test–retest reliability of resting-state fMRI metrics and optimization of scan length. Front. Neurosci. 16, (2022).

66. Wang, P. et al. Amplitude of low-frequency fluctuation (ALFF) may be associated with cognitive impairment in schizophrenia: a correlation study. BMC Psychiatry 19, 30 (2019).

67. Song, C., Boly, M., Tagliazucchi, E., Laufs, H. & Tononi, G. fMRI spectral signatures of sleep. Proc. Natl. Acad. Sci. 119, e2016732119 (2022).

68. Fukunaga, M. et al. Large-amplitude, spatially correlated fluctuations in BOLD fMRI signals during extended rest and early sleep stages. Magn. Reson. Imaging 24, 979–992 (2006).

69. Horovitz, S. G. et al. Low frequency BOLD fluctuations during resting wakefulness and light sleep: A simultaneous EEG-fMRI study. Hum. Brain Mapp. 29, 671–682 (2008).

70. Bohr, T. et al. The glymphatic system: Current understanding and modeling. iScience 25, 104987 (2022).

71. Persson, N. D. Å., Uusalo, P., Nedergaard, M., Lohela, T. J. & Lilius, T. O. Could dexmedetomidine be repurposed as a glymphatic enhancer? Trends Pharmacol. Sci. 43, 1030–1040 (2022).

72. Lilius, T. O. et al. Dexmedetomidine enhances glymphatic brain delivery of intrathecally administered drugs. J. Controlled Release 304, 29–38 (2019).

73. Benveniste, H. et al. Anesthesia with Dexmedetomidine and Low-dose Isoflurane Increases Solute Transport via the Glymphatic Pathway in Rat Brain When Compared with High-dose Isoflurane. Anesthesiology 127, 976–988 (2017).

74. Wang, S. et al. Dexmedetomidine improves the circulatory dysfunction of the glymphatic system induced by sevoflurane through the PI3K/AKT/ΔFosB/AQP4 pathway in young mice. Cell Death Dis. 15, 448 (2024).

75. Liu, X. et al. Propofol attenuates low-frequency fluctuations of resting-state fMRI BOLD signal in the anterior frontal cortex upon loss of consciousness. NeuroImage 147, 295–301 (2017).

76. Hillman, E. M. C. Coupling Mechanism and Significance of the BOLD Signal: A Status Report. Annu. Rev. Neurosci. 37, 161–181 (2014).

77. Kim, S.-G. & Ogawa, S. Biophysical and physiological origins of blood oxygenation level-dependent fMRI signals. J. Cereb. Blood Flow Metab. 32, 1188–1206 (2012).

78. Pauling, L. & Coryell, C. D. The Magnetic Properties and Structure of Hemoglobin, Oxyhemoglobin and Carbonmonoxyhemoglobin. Proc. Natl. Acad. Sci. 22, 210–216 (1936).

79. McAvoy, M. P., Tagliazucchi, E., Laufs, H. & Raichle, M. E. Human non-REM sleep and the mean global BOLD signal. J. Cereb. Blood Flow Metab. 39, 2210–2222 (2019).

80. Deng, S. et al. Hemodynamic and metabolic correspondence of resting-state voxel-based physiological metrics in healthy adults. NeuroImage 250, 118923 (2022).

81. Wasilczuk, A. Z., et al. Hormonal basis of sex differences in anesthetic sensitivity. Proc. Natl. Acad. Sci. 121, e2312913120 (2024).

82. Lennertz, R., et al. Connected consciousness after tracheal intubation in young adults: an international multicentre cohort study. BJA Br. J. Anaesth. 130, e217–e224 (2023).

83. Braithwaite, H. E. et al. Impact of female sex on anaesthetic awareness, depth, and emergence: a systematic review and meta-analysis. Br. J. Anaesth. 131, 510–522 (2023).

84. Li, Y.-Y., Ge, D.-J., Li, J.-Y. & Qi, B. Sex Differences in the Morphine-Sparing Effects of Intraoperative Dexmedetomidine in Patient-Controlled Analgesia Following General Anesthesia: A Consort-Prospective, Randomized, Controlled Clinical Trial. Medicine (Baltimore) 95, e3619 (2016).

85. Buckner, R. L. et al. Cortical Hubs Revealed by Intrinsic Functional Connectivity: Mapping, Assessment of Stability, and Relation to Alzheimer’s Disease. J. Neurosci. 29, 1860–1873 (2009).

86. Crossley, N. A. et al. The hubs of the human connectome are generally implicated in the anatomy of brain disorders. Brain 137, 2382–2395 (2014).

87. Tononi, G. & Cirelli, C. Sleep and the Price of Plasticity: From Synaptic and Cellular Homeostasis to Memory Consolidation and Integration. Neuron 81, 12–34 (2014).

88. Jenkinson, M., Beckmann, C. F., Behrens, T. E. J., Woolrich, M. W. & Smith, S. M. FSL. NeuroImage 62, 782–790 (2012).

89. Makris, N. et al. Decreased volume of left and total anterior insular lobule in schizophrenia. Schizophr. Res. 83, 155–171 (2006).

90. Frazier, J. A. et al. Structural Brain Magnetic Resonance Imaging of Limbic and Thalamic Volumes in Pediatric Bipolar Disorder. Am. J. Psychiatry 162, 1256–1265 (2005).

91. Desikan, R. S. et al. An automated labeling system for subdividing the human cerebral cortex on MRI scans into gyral based regions of interest. NeuroImage 31, 968–980 (2006).

92. Goldstein, J. M. et al. Hypothalamic Abnormalities in Schizophrenia: Sex Effects and Genetic Vulnerability. Biol. Psychiatry 61, 935–945 (2007).

93. Tzourio-Mazoyer, N. et al. Automated Anatomical Labeling of Activations in SPM Using a Macroscopic Anatomical Parcellation of the MNI MRI Single-Subject Brain. NeuroImage 15, 273–289 (2002).

94. Tournier, J.-D. et al. MRtrix3: A fast, flexible and open software framework for medical image processing and visualisation. NeuroImage 202, 116137 (2019).

95. Smith, R. E., Tournier, J.-D., Calamante, F. & Connelly, A. SIFT2: Enabling dense quantitative assessment of brain white matter connectivity using streamlines tractography. NeuroImage 119, 338–351 (2015).

96. Statistical Parametric Mapping: The Analysis of Functional Brain Images. (Elsevier, 2011).

97. Chen, S. et al. Global Functional Connectivity at Rest Is Associated with Attention: An Arterial Spin Labeling Study. Brain Sci. 13, 228 (2023).

98. Mahadevan, A. S., Tooley, U. A., Bertolero, M. A., Mackey, A. P. & Bassett, D. S. Evaluating the sensitivity of functional connectivity measures to motion artifact in resting-state fMRI data. NeuroImage 241, 118408 (2021).

99. James, O., Park, H. & Kim, S.-G. Impact of sampling rate on statistical significance for single subject fMRI connectivity analysis. Hum. Brain Mapp. 40, 3321–3337 (2019).

100. Whitfield-Gabrieli, S. & Nieto-Castanon, A. Conn: a functional connectivity toolbox for correlated and anticorrelated brain networks. Brain Connect. 2, 125–141 (2012).

101. Nieto-Castanon, A. Handbook of fcMRI Methods in CONN. (Hilbert Press, Boston, MA, 2020).

102. Yang, H. et al. Amplitude of low frequency fluctuation within visual areas revealed by resting-state functional MRI. NeuroImage 36, 144–152 (2007).

103. Alsop, D. C. et al. Recommended implementation of arterial spin-labeled perfusion MRI for clinical applications: A consensus of the ISMRM perfusion study group and the European consortium for ASL in dementia. Magn. Reson. Med. 73, 102–116 (2015).

104. Weinstein, S. M. et al. A simple permutation-based test of intermodal correspondence. Hum. Brain Mapp. 42, 5175 (2021).

105. Mitchell, S. M., Lange, S. & Brus, H. Gendered Citation Patterns in International Relations Journals. Int. Stud. Perspect. 14, 485–492 (2013).

106. Dion, M. L., Sumner, J. L. & Mitchell, S. M. Gendered Citation Patterns across Political Science and Social Science Methodology Fields. Polit. Anal. 26, 312–327 (2018).

107. Caplar, N., Tacchella, S. & Birrer, S. Quantitative evaluation of gender bias in astronomical publications from citation counts. Nat. Astron. 1, 1–5 (2017).

108. Maliniak, D., Powers, R. & Walter, B. F. The Gender Citation Gap in International Relations. Int. Organ. 67, 889–922 (2013).

109. Dworkin, J. D. et al. The extent and drivers of gender imbalance in neuroscience reference lists. Nat. Neurosci. 23, 918–926 (2020).

110. Smith, S. M. & Nichols, T. E. Threshold-free cluster enhancement: Addressing problems of smoothing, threshold dependence and localisation in cluster inference. NeuroImage 44, 83–98 (2009).

